# Cerebrospinal fluid reference proteins increase accuracy and interpretability of biomarkers for brain diseases

**DOI:** 10.1101/2023.06.08.544222

**Authors:** Linda Karlsson, Jacob Vogel, Ida Arvidsson, Kalle Åström, Shorena Janelidze, Kaj Blennow, Sebastian Palmqvist, Erik Stomrud, Niklas Mattsson-Carlgren, Oskar Hansson

**Author notes:** Correspondence to: Linda Karlsson Full address: Biomedical Centre, C11, SE-22184 Lund, Sweden. or Oskar Hansson Full address: Memory Clinic, Skåne University Hospital, SE-20502 Malmö, Sweden.

## Abstract

Cerebrospinal fluid (CSF) biomarkers reflect brain pathophysiology and are used extensively in translational research as well as in clinical practice for diagnosis of neurological diseases, e.g., Alzheimer’s disease (AD). However, CSF biomarker concentrations may be influenced by non-disease related mechanisms which vary between individuals, such as CSF production and clearance rates. Here we use a data-driven approach to demonstrate the existence of inter-individual variability in mean CSF protein levels. We show that these non-disease related differences cause many commonly reported CSF biomarkers to be highly correlated, thereby producing misleading results if not accounted for. To adjust for this inter-individual variability, we identified and evaluated high-performing reference proteins which improved the diagnostic accuracy of key CSF AD biomarkers. Our novel reference protein method attenuates the risk for false positive findings, and improves the sensitivity and specificity of CSF biomarkers, with broad implications for both research and clinical practice.

## Introduction

Neurodegenerative disorders and dementia are common and have increasing prevalence world-wide.^1^ The need for precise and reliable diagnostic techniques to identify, examine and monitor these diseases is growing. One informative and cost-effective diagnostic technique is the measurement of protein concentrations in cerebrospinal fluid (CSF), here referred to as CSF biomarkers.^2, 3^ In Alzheimer’s disease (AD), which is the most common neurodegenerative disease, CSF biomarkers are used in clinical practice as diagnostic tools.^2^ Neuropathologically, AD is defined by the combined presence of amyloid(A)-β plaques and tau-neurofibrillary tangles. CSF biomarkers related to these pathologies include Aβ42 and soluble phosphorylated(P)-tau.^4^ These CSF markers can substantially improve the diagnostic work-up of the disease, which is becoming increasingly important due to recent development of effective disease modifying-treatments for AD.^5–7^ However, the use of CSF biomarkers may be complicated by inter-individual variability in certain physiological phenomena, such as rates of CSF production, and rates of CSF clearance.^8–10^ Such inter-individual differences could lead to differences in mean CSF protein levels^11^, which could impact the overall performance of CSF biomarkers.^12–14^ Hypothetically, adjustment for individual mean CSF protein levels could optimize the performance of already efficient CSF biomarkers, reduce false positive findings (by attenuating biomarker associations that are driven by the mean CSF protein level), and increase the likelihood of making new biologically and clinically relevant discoveries.

In AD research and clinical practice, CSF Aβ42 and P-tau, together with a CSF biomarker of neuronal injury (i.e., CSF total-tau or neurofilament light chain), can be used for AT(N) (amyloid, tau, neurodegeneration) *in vivo* classification of AD pathology.^15, 16^ This system makes it possible to categorize a person as biomarker positive or negative, where low CSF Aβ42 levels indicate Aβ plaque pathology (“A”) and high CSF P-tau181 levels indicate tau tangle pathology (“T”).^17–23^ AT(N) grouping is an effective way to differentiate individuals without AD (A-/T-) from those with AD (A+T+). However, many studies have reported findings in the group with isolated P-tau pathology (A-T+; i.e., both high Aβ42 and P-tau181), which is more controversial.^15, 24–26^ It is unclear if this A-T+ definition is biologically relevant or mainly a result of inter-individual differences in mean CSF levels (leading to more concentrated CSF in some individuals with higher levels of both Aβ42 and P-tau181).

Besides well-established CSF biomarkers used in clinical practice, CSF proteins are often studied to understand underlying disease mechanisms in humans affected by AD or other neurodegenerative diseases. In such studies, it has been suggested that CSF levels of many microglia-related proteins (like sTREM2 or TAM receptors [sAXL and sTYRO3]) are strongly correlated with CSF P-tau181 and increased not only in A+T+ individuals, but also in A-T+ individuals, linking these neuroinflammatory changes more to P-tau pathology than Aβ.^18, 19^ In addition, other CSF markers have been seen to correlate with CSF P-tau181 levels. For example, we and others have reported that the astrocytic biomarker YKL-40 and the Parkinson’s disease-related biomarker α-synuclein are strongly associated with P-tau181 in CSF, which was interpreted as that these brain pathological changes co-vary.^27–29^ It is unclear whether such findings are mainly driven by inter-individual differences in mean CSF protein levels, or remain robust when accounting for this property. Moreover, the impact of mean CSF levels might also be of importance in proteomic studies, when identifying subpopulations with different CSF expression profiles^30^, or in genome-wide protein quantitative trait loci (pQTL) studies, looking at associations between genetic variants and protein levels.^14^

One striking example that highlights the potential of adjusting for processes related to CSF dynamics in the context of AD CSF biomarkers exists. CSF Aβ42 shows improved concordance with amyloid positron emission tomography (PET, a well-established neuroimaging method to make aggregated brain amyloid in AD visible) when normalized for CSF Aβ40 levels, where the latter is not affected by the disease process.^31, 32^ Aβ40 is closely linked to Aβ42 since both peptides come from the same proteolytic pathway^33^, but Aβ40 may also partly represent an individual’s mean CSF protein level and could potentially improve performance of other biomarkers as well. This idea has been tested for CSF P-tau181, where the results suggested that the diagnostic accuracy improved when adjusting for inter-individual differences in CSF Aβ40 levels.^13^ In order to examine Aβ40’s generalizability as a reference protein, it needs to be further evaluated. In addition to Aβ40, other efficient CSF reference proteins may exist that can improve the clinical performance of key CSF biomarkers.

Consequently, our overarching aim was to establish the concept of inter-individual differences in mean CSF protein levels, and to search for optimal reference protein candidates that could be used to account for this CSF dynamic in a robust way. We analyzed 2,944 CSF proteins (including CSF Aβ40) from 830 participants in a data-driven manner. We hypothesized that adjusting for certain reference proteins could improve the diagnostic accuracy of AD CSF biomarkers, and we evaluated this across a range of outcome measures, biomarkers, and AD cohorts. We also hypothesized that several previously reported CSF biomarker findings would be altered or attenuated when biomarkers were normalized to reference proteins. Specifically, we studied whether several strong and recognized correlations of CSF protein concentrations remained robust when normalizing to the identified reference proteins, both in relation to each other and to genetic variants.

## Results

The study included 830 participants from the Swedish BioFINDER-2 (BF2) cohort and 904 participants from Swedish BioFINDER-1 BF1 cohort, all with complete OLINK CSF protein and CSF Aβ40 concentration measures (2,944 in BF2 and 369 in BF1). BF2 was randomly split into a training set (80%, n=658) and test set (20%, n=172). Throughout this work, the training dataset of BF2 was used for all exploratory work. The BF2 test dataset was used to evaluate findings, and BF1 was used for external validation. To find and assess appropriate reference proteins, their performance was evaluated in three logistic regression models. The first model predicted tau-PET (a well-established neuroimaging method to demonstrate fibrillary tau deposition in AD) positivity with CSF P-tau181 (**P-tau181→TauPET**). The second predicted Aβ-PET (a well-established neuroimaging method to demonstrate fibrillary amyloid deposition in AD) positivity with CSF Aβ42 (**Aβ42→AβPET**). The third predicted future conversion to AD dementia with CSF P-tau181 (**P-tau181→ADDconv**). The first two models were used to search for suitable reference proteins while the third was used for validation of reference proteins. A flowchart and details of the complete reference search/evaluation pipeline, together with all data splitting details and demographics, are presented in Fig. 1 and Tab. 1.

**FIGURE 1:**
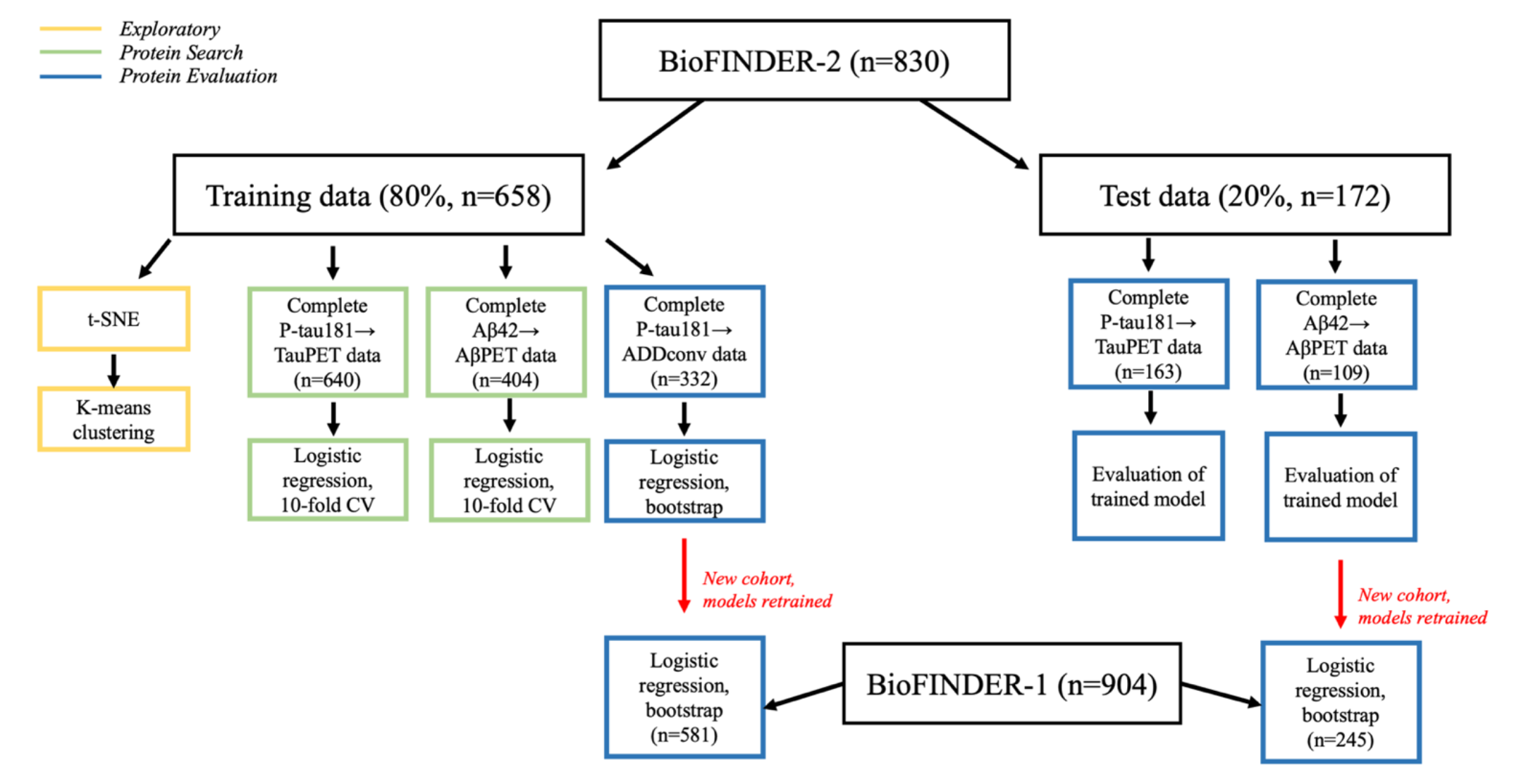
Flowchart of reference protein search and evaluation pipeline. BF2 was filtered by participants with CSF OLINK and CSF Aβ40 measurements (n=830). Thereafter, the dataset was split into 80% training (n=658) and 20% testing (n=172). During the exploratory phase, all BF2 training data was used. Next, the two models **P-tau181→TauPET** and **Aβ42→AβPET** were used to search for reference proteins in the protein search phase. The proposed candidates were evaluated in corresponding models for unseen test data, and in a new third model **P-tau181→ADDconv** on the training data. The findings were further validated in the independent cohort BF1 (n=904) for the two models **Aβ42→AβPET** and **P-tau181→ADDconv**. The model **P-tau181→TauPET** was not evaluated in BF1 as baseline tau-PET data did not exist. Complete data refers to no missing values for any of the relevant variables and was a filtering step in all models.

**TABLE 1:**
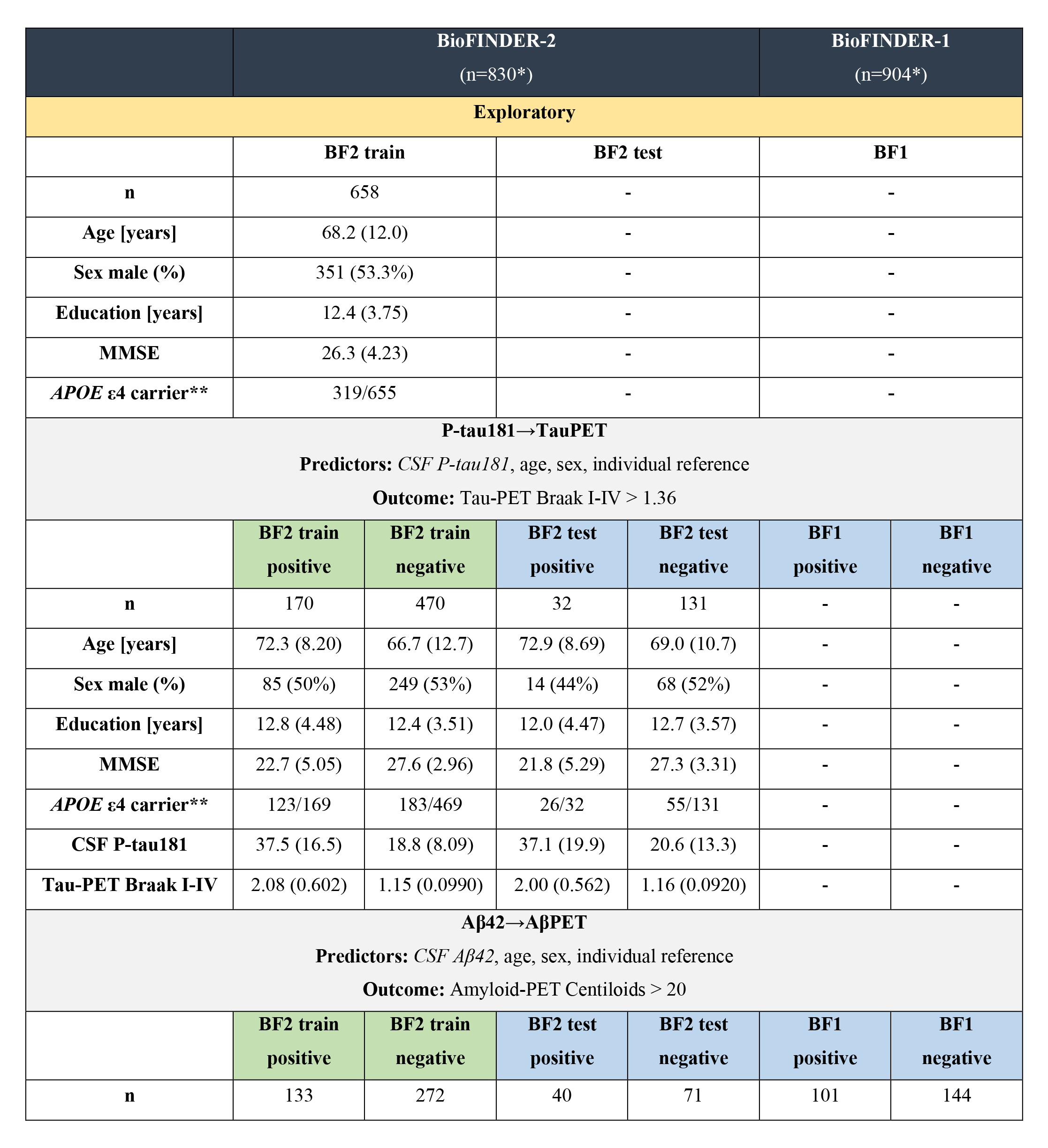
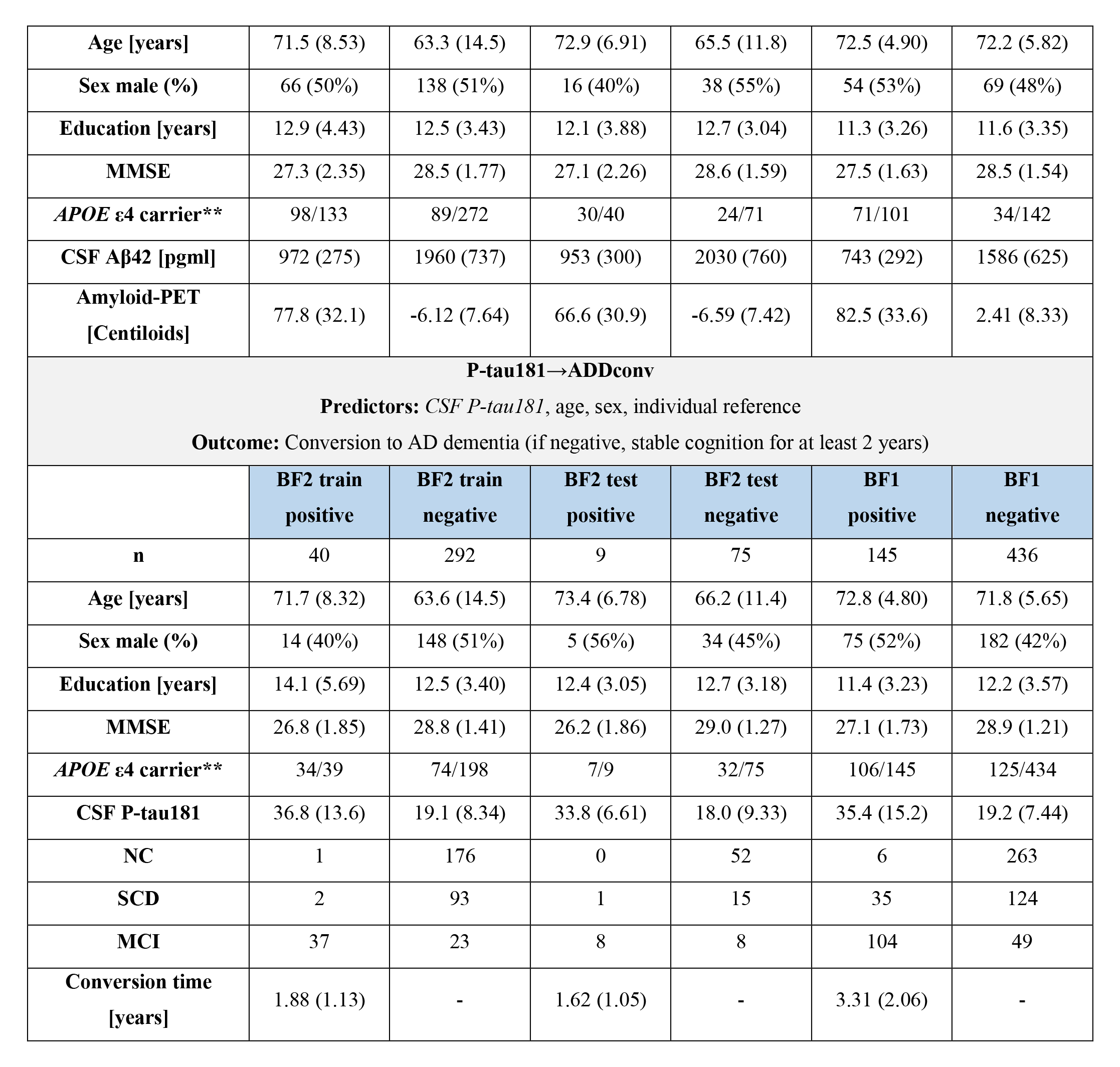
Demographics for all data paths in Fig. 1. Details of all data used for exploration (yellow), protein search (green) and protein evaluation (blue) in concordance with the different data paths in Fig. 1. The regression models (gray) all include age, sex and individual reference, but differ by main predictor (italics) and outcome. The differences generated a variation in number of participants and demographics for each model, depending on the data available. Exploratory work was only performed on training data in BF2. In BF1, tau-PET data did not exist. Abbreviations: mini mental state examination (MMSE), positron emission tomography (PET), normal cognition (NC), subjective cognitive decline (SCD), mild cognitive impairment (MCI).

### Many CSF proteins vary in concordance with the mean CSF protein level

We sorted the 2,944 standardized CSF protein concentrations according to their associations with the individual mean CSF protein level (Supplementary Fig. 1) and visualized the results in Fig. 2. Fig 2a indicates that, within a random subsample of individuals, there are several participants that systematically have high or low values across several hundred proteins. This phenomenon is evident across the full training dataset of 658 participants (Fig 2b), where nearly half of all proteins measured appear to show highly consistent individual variation. When removing proteins of low detectability, this pattern becomes even clearer (Supplementary Fig. 2), emphasizing that most proteins that are highly expressed in CSF (and therefore likely to be nominated in CSF biomarker studies) vary in concordance with the mean CSF protein level. The AD CSF biomarkers P-tau181 (β=0.34, *P*<1e-20) and Aβ42 (β=0.24, *P*<1e-10) were strongly associated with the mean CSF protein level (Fig. 2c). As expected, Aβ40 (β=0.44, *P*<1e-37) showed a strong association with the mean CSF protein level (Fig. 2c).

**FIGURE 2:**
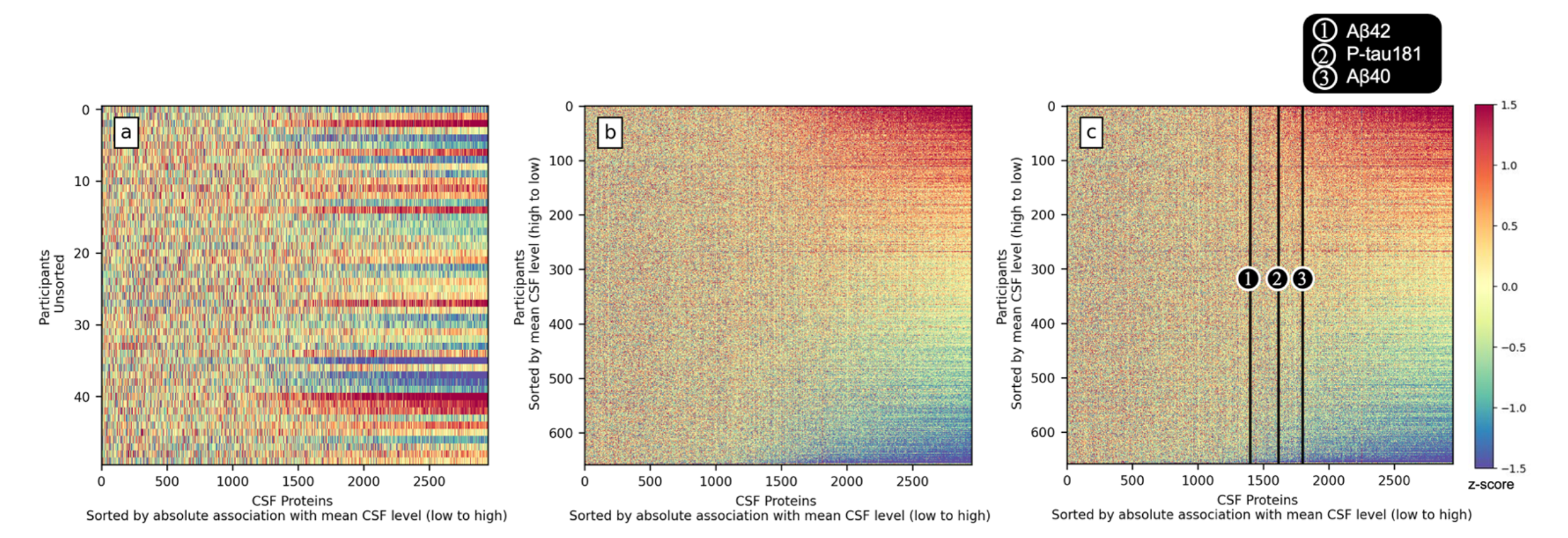
Many CSF proteins vary in concordance with an individual protein level. For each participant (row), the standardized concentration of 2,944 CSF proteins, sorted by increasing absolute association with the mean CSF protein level (Supplementary Fig. 1), is displayed. Systematic blue horizontal lines can be seen for individuals with consistently low values across most proteins, and correspondingly red horizontal lines for individuals with high values across most proteins (all relative to the total sample). **a)** a subset of 50 randomly selected participants. **b)** all 658 participants sorted by mean CSF level. **c)** same as b) but also including biomarkers Aβ42 and P-tau181, which together with Aβ40 are marked out. The further right the protein is located, the more associated with the mean CSF level and therefore more strongly confounded by the mean CSF protein level when used as a biomarker.

To further understand potential underlying mechanisms of mean CSF protein level differences, we investigated the mean CSF protein level’s association with age, sex, education level, intracranial volume, gray matter volume and ventricular volume. In a multiple linear regression model, significant associations with a higher mean CSF protein level were found for higher age (β=0.544, *P*=4e-31), male sex (β=-0.159, *P*=2e-4), and lower ventricular volume (β=-0.321, *P*=5e-11), see Supplementary Tab. 1. Similar results were seen when evaluating Aβ- negative cognitively normal participants only.

### A cluster with superior CSF reference protein qualities

We next examined clustering of the CSF protein concentrations to identify proteins of similar characteristics. We used t-SNE dimensionality reduction^34^, applied to the high dimensional space of 658 participants (Fig. 3a). As the algorithm optimizes to preserve similarity of pairwise points, proteins of short distance in Fig. 3a can be interpreted as similarly expressed. In Fig. 3c-3h, clustering characteristics of the t-SNE space are compared against several metrics relevant in search of reference proteins. A reference protein should be associated with the mean CSF protein level (Fig. 3c). The mean CSF protein level was highly associated with ventricular volume when adjusting for age and sex, suggesting that the mean CSF level is partly driven by dilution. Therefore, lower levels of optimal reference proteins could be associated with larger volume when adjusting for age and sex, suggesting that the mean CSF level is partly driven by dilution. Therefore, lower levels of optimal reference proteins could be associated with larger ventricular volumes (Fig. 3d). A potential reference protein for a given model predictor, that can perform better than simply using the mean CSF level of all proteins, should co-vary with the main predictor during normal physiology but not in disease. Therefore, an AD reference protein should likely have a high correlation with key biomarkers like P-tau181 and Aβ42 in cognitively unimpaired Aβ-negative participants, which are not considered to have the disease (Fig. 3e and 3f). Lastly, a well-performing reference protein should improve the predictive performance of key biomarkers, which was evaluated by comparing results of i) using P-tau181 together with a potential reference protein to predict tau-PET outcome (**P-tau181→TauPET)** or ii) using Aβ42 together with a potential reference protein to predict Aβ-PET outcome **(Aβ42→AβPET)** (Fig. 3g and 3h).

**FIGURE 3:**
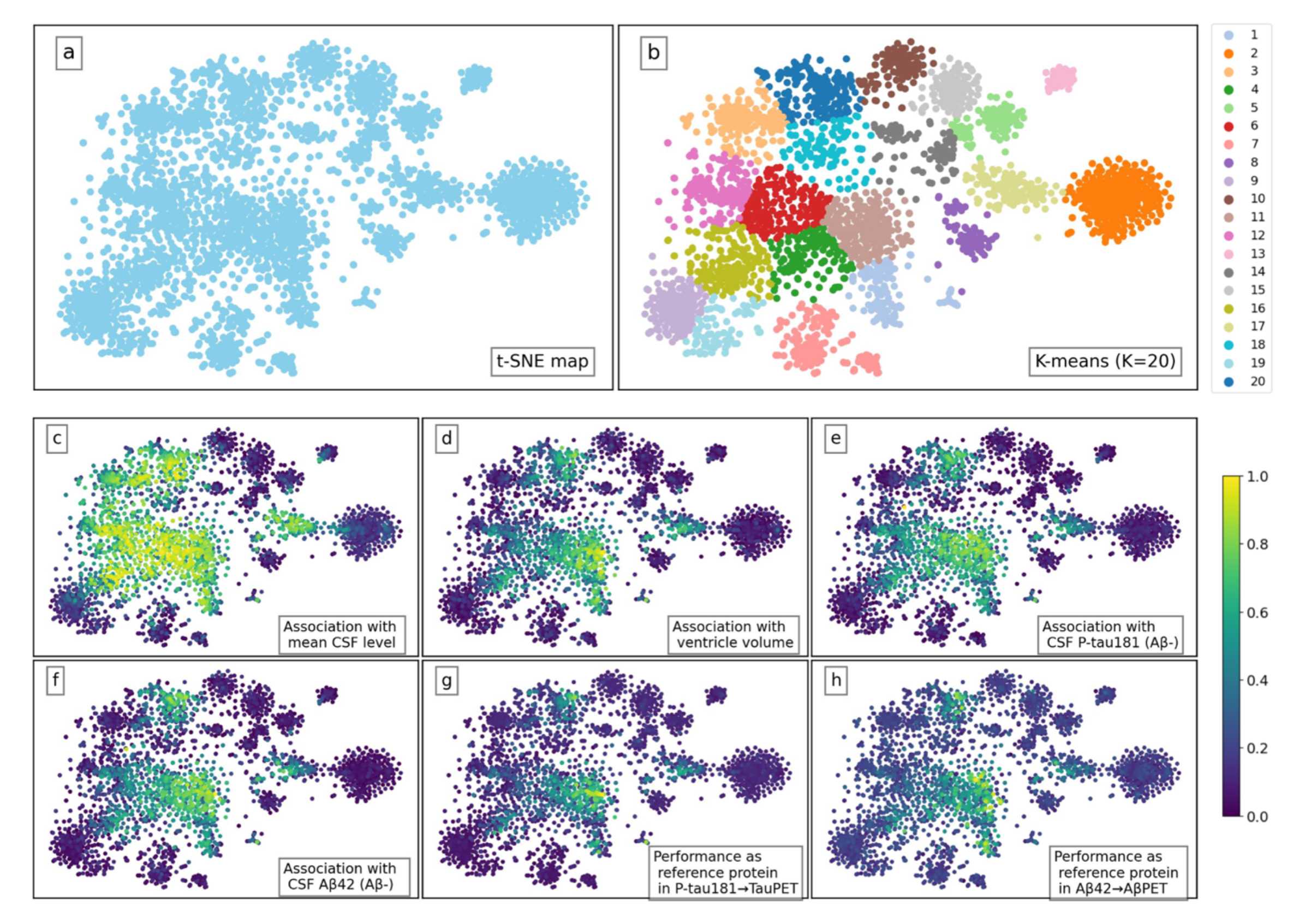
Dimensionality reduction reveals a cluster of CSF proteins with desired reference protein characteristics. T-distributed stochastic neighbor embedding (t-SNE), reducing the high dimensional space of 658 participants to a two-dimensional one. Each scatter point illustrates one of the 2,944 CSF proteins, with relative similarity of pairwise proteins aimed to be preserved. In c)-h), min-max scaling for six different criteria has been performed to visualize relative differences within the space, all plots ranging between 0-1 (dark blue to yellow). In c)-f) associations are analyzed as absolute β-coefficients, all adjusted for age and sex. **a)** The raw t-SNE map. **b)** Semi-supervised K-means (K=20) clustering of t-SNE map, aiming to separate the evident clusters from t-SNE dimensionality reduction and areas of overlap in c)-h). t-SNE map colored by **c)** absolute association with mean CSF level; **d)** absolute association with ventricle volume; **e)** absolute association with biomarker CSF P-tau181 (cognitively unimpaired Aβ-negative participants only); **f)** absolute association with biomarker CSF Aβ42 (cognitively unimpaired Aβ-negative participants only); **g)** model performance when used as reference protein in **P-tau181→TauPET**; **h)** model performance when used as reference protein in **Aβ42→AβPET**.

A semi-supervised K-means (K=20) clustering algorithm^35^ was utilized to divide the t-SNE space and identify a subset of potential reference proteins. The supervision was performed by selecting K and the random initialization so that clear structural clusters were separated and areas of overlap in Fig. 3c-3h could be examined in more detail. The resulting K-means clustering can be seen in Fig. 3b. The clustering was relatively well in line with the OLINK panel division (see Supplementary Fig. 3), where the area of interest was likely to not benefit a certain panel.

The AUC of each cluster in the two regression models **P-tau181→TauPET** and **Aβ42→AβPET** were evaluated in Fig. 4a and 4b. The AUCs without using an individual reference were 0.865 and 0.934 respectively. By analyzing the performance cluster-wise, we aimed to target protein expression characteristics rather than single findings and hence remove top performances biased by data. As seen in both Fig. 4a and 4b, and as expected from the overlapping areas in Fig. 3, cluster 11 (n_proteins_=219) stands out as the best performing cluster (mean AUC ± std: 0.896 ± 0.0187 and 0.947 ± 0.00698 for **P-tau181→TauPET** and **Aβ42→AβPET** respectively).

**FIGURE 4:**
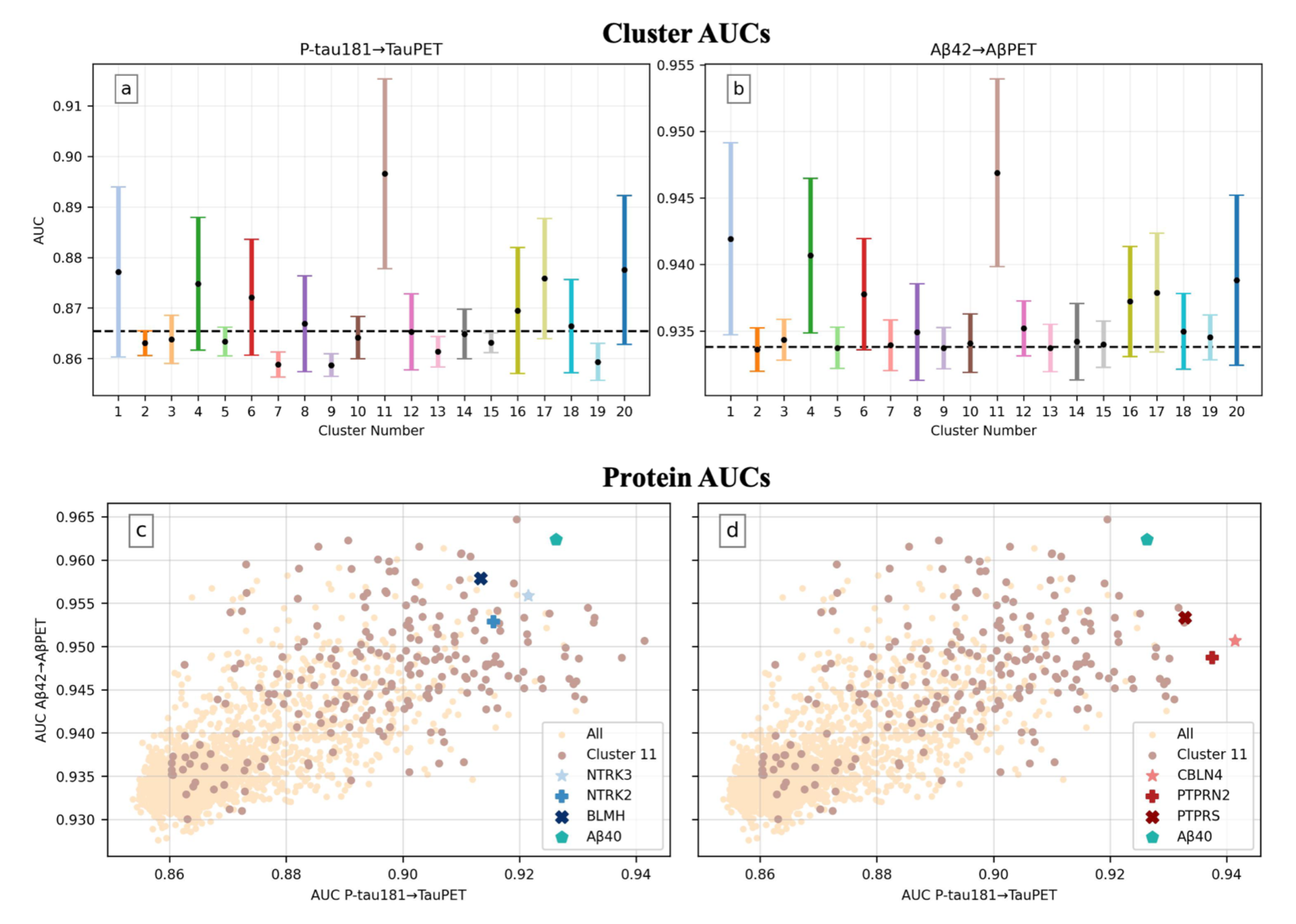
Cluster 11 stands out as superior when used as reference in models P-tau181→TauPET and Aβ42→AβPET. In **a)** and **b)** each error bar represents the cluster’s mean performance ± one standard deviation when the proteins of the cluster are used as a participant’s individual reference (one protein at a time, all proteins evaluated once, adjusting for age and sex). The dashed lines correspond to the models’ AUCs without using a reference (0.865 in a) and 0.934 in b)). Each cluster is colored as in Fig. 3b. For both models, cluster 11 stands out as the best performing cluster on average. In **c)** and **d)** every scatter point corresponds to the result when evaluating that protein as individual reference. Consequently, the top right corner contains proteins of most interest. Proteins in cluster 11, Aβ40 and three other reference protein candidates are highlighted. Note that **c)** and **d)** are identical apart from the highlighted markers. **a)** results for **P-tau181→TauPET** by cluster. **b)** results for **Aβ42→AβPET** by cluster**. c)** proteins NTRK3, NTRK2 and BLMH selected as general reference proteins, after enforcing the **selection criteria**. **d)** proteins CBLN4, PTPRN2 and PTPRS selected as P-tau181 specific reference proteins, as they improved **P-tau181→TauPET** the most. All models were evaluated using 10-fold-cross-validation on the BF2 training dataset.

We performed cell type expression and cellular component pathway enrichment analyses on cluster 11 to further investigate the characteristics of promising reference proteins from a biological perspective. Cluster 11 had high expression in mainly neuronal cells (Supplementary Fig. 4) but showed no significant expression difference compared to the other 2,725 OLINK proteins, which was assessed with a bootstrap enrichment test. Cluster 11 was enriched on cell surfaces and membranes (Supplementary Fig. 5).

### Identification of general and biomarker-specific reference proteins

To further validate single robust reference proteins, specific candidates from cluster 11 that resulted in top AUC scores for the two models **P-tau181→TauPET** and **Aβ42→AβPET** were identified (Fig. 4c and 4d). While this extensive dataset of 2,944 proteins allowed for great exploration possibilities, it also limited the validation opportunities in other cohorts. Additionally, some cohort-specific biases in our data were still expected, even after adding robustness by only looking at the subset of proteins from cluster 11. We hence did not expect small AUC differences between single proteins to be significant. Taking these factors into account, we selected three reference protein candidates in addition to Aβ40 (also in cluster 11), for further examination, based on the following **selection criteria**:

1. The protein was in cluster 11.
2. The inclusion of the protein led to an increased AUC score for both the **P-tau181→TauPET** and **Aβ42→AβPET** model.
3. The protein was measured in our independent validation cohort (BF1).

The resulting three proteins, NTRK3, NTRK2 and BLMH, are marked out in Fig. 4c. See *Supplementary Reference Candidates* for a biological description of the three proteins and BF2 data information further confirming the proteins’ potential as suitable references (e.g., high association with mean CSF level, small concentration differences between diagnostic groups, high association with the main predictors and low model performance when used without a main predictor).

We hypothesized that each main predictor may have one or several *optimal* reference candidates. An optimal reference is co-varying to a higher extent with the main predictor during normal physiology but not in disease. For Aβ42, a natural biomarker specific reference is Aβ40, as both are generated from the amyloid precursor protein (APP)^33^ and hence largely follow the same biological pathway. To further evaluate this concept of biomarker-specific reference proteins, we investigated the possibility of finding exceptionally high-performing references for CSF P-tau181. There is high relevance in identifying optimal references for CSF P-tau181, as there is no state-of-the-art way of normalizing this biomarker. Furthermore, as CSF P-tau181 was more correlated with the mean CSF level than Aβ42, it may have more potential for improvement when adjusting for a reference protein. This idea was strengthened by the results shown in Fig. 4, where the AUC improvement was considerably larger for **P-tau181→TauPET** than **Aβ42→AβPET** when adjusting for a reference (max AUC improvement: 0.076 versus 0.031, cluster 11 mean AUC improvement: 0.031 versus 0.013). We therefore identified the three proteins in cluster 11 that improved **P-tau181→TauPET** the most: CBLN4, PTPRN2 and PTPRS in Fig. 4d. See *Supplementary Reference Candidates* for a biological description of the three proteins and BF2 data information further confirming the proteins’ potential as suitable references.

### Adjusting for reference proteins improves biomarker performance

In accordance with the flowchart in Fig. 1, five combinations of models and datasets were evaluated. For all models, performance was compared between no reference and Aβ40, mean CSF level and the three general reference protein candidates NTRK3, NTRK2, BLMH as reference. Additionally, as the three candidates were highly correlated (Pearson correlation 0.85-0.91, see Supplementary Fig. 6a), the first component of a singular value decomposition was also evaluated as a possible reference, created from the three candidates (SVD1). For all models using BF2 data, the P-tau181-specific reference protein candidates (CBLN4, PTPRN2 and PTPRS, Pearson correlation 0.79-0.89, see Supplementary Fig. 6b) and their corresponding first component of a singular value decomposition (SVD2) were evaluated as well.

The two models **P-tau181→TauPET** and **Aβ42→AβPET** were retrained on the full BF2 training dataset and evaluated on the BF2 test dataset (Fig. 5a-5d and Supplementary Tab. 2). For both models and all references, the performance significantly increased (AUC = 0.895-0.946 and 0.977-0.992, *P*<0.05, for **P-tau181→TauPET** and **Aβ42→AβPET** respectively) compared to no reference (AUCs = 0.828 and 0.966). Additionally, using a single reference protein or an SVD of three candidates (AUCs = 0.908-0.946 and 0.982-0.992) also outperformed using the mean CSF level as reference (AUCs = 0.895 and 0.977). For **P-tau181→TauPET**, top performance was reached when adjusting for SVD2 (AUC = 0.946, *P*<0.01), closely followed by CBLN4 (AUC = 0.944, *P*<0.01). For **Aβ42→AβPET**, top performance was reached when adjusting for Aβ40 (AUC = 0.992, *P*<0.05).

**FIGURE 5:**
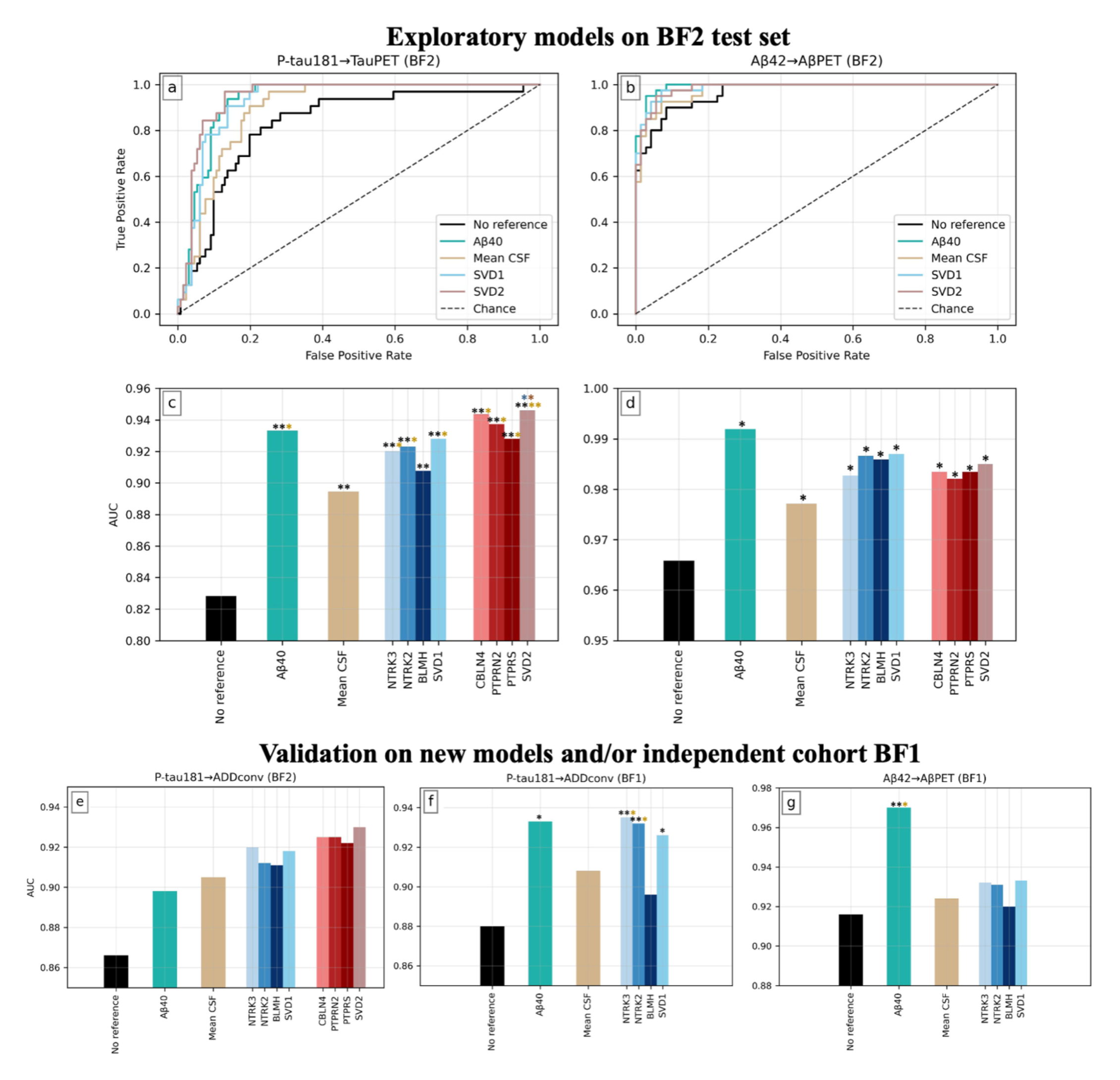
Performance evaluation of all models with and without references. ROC curves in **a)** and **b),** and corresponding AUCs in **c)** and **d)**, for the two models **P-tau181→TauPET** and **Aβ42→AβPET**, evaluated on the BF2 test dataset. AUCs for **e)** model **P-tau181→ADDconv** on the training dataset BF2, **f)** model **P-tau181→ADDconv** on BF1 and **g)** model **Aβ42→AβPET** on BF1. No reference corresponds to the model without use of a reference protein, consistently outperformed by all tested references: Aβ40, mean CSF level, NTRK3, NTRK2, BLMH, SVD1, CBLN4, PTPRN2, PTPRS and SVD2. Quantitative details can be found in Supplementary Tab. 2. For visualization purposes, the ROC curves do not show all protein candidates but solely the corresponding SVDs. \**P* < 0.05, \*\**P* < 0.01 compared to the bar of same color as asterisk.

To validate the generalizability of the reference candidates, **Aβ42→AβPET** was applied in BF1 and the new model **P-tau181→ADDconv** was applied in both BF2 and BF1 (Fig. 5e- 5g and Supplementary Tab. 2). Note that measurements of CBLN4, PTPRN2 and PTPRS were not available in BF1. Again, using no reference consistently resulted in the lowest performance (AUCs = 0.866, 0.880 and 0.916 for **P-tau181→ADDconv** in BF2, **P-tau181→ADDconv** in BF1, and **Aβ42→AβPET** in BF1 respectively). For **P-tau181→ADDconv** in BF2 (Fig. 5e), no significant improvements were achieved, most likely due to the small sample size (n_pos_=40, n_neg_=292). However, the same trends as in Fig. 5c were seen, where the proposed P-tau181- specific reference candidates again achieved top performance, together with corresponding SVD2 (AUCs = 0.922-0.930). For the same model **P-tau181→ADDconv** in BF1 (Fig. 5f), the available data was larger (n_pos_=145, n_neg_=436), and several significant improvements were achieved both compared to no reference and mean CSF level as reference, with NTRK3 showing best performance (AUC = 0.935, *P*<0.01). For **Aβ42→AβPET** in BF1 (Fig. 5g), a significant improvement was only achieved with Aβ40 (AUC=0.970, *P*<0.05), which was clearly superior to all other tested references.

### Reference proteins explain discordance between CSF and PET tau positivity

We investigated how adjusting for an individual reference protein affected concordance between AT(N) grouping for CSF P-tau181 and tau-PET (Fig. 6). For this analysis, all participants from the BF2 training dataset with tau-PET data were included (n=640). A-grouping was performed with CSF Aβ42/Aβ40 (cutoff 0.08^36^). T-grouping was made using 1) no reference (Fig. 6a, CSF cutoff of P-tau181>21.8 pg/ml, as in e.g. Suárez-Calvet, M. et al^18^) and 2) the reference protein candidate CBLN4 (Fig. 6c, CSF cutoff of P-tau181 > 39.0 + 10.1c_CBLN4_, adapted from a logistic regression model). In addition, the grouping methods were compared against 3) tau-PET grouping (Fig. 6b and Fig. 6d). The concordance between CSF and PET grouping increased both visually and quantitively when not requiring a cutoff based on CSF P-tau181 only, but also accounting for the reference protein. The accuracy increased from 76% to 89% (Fig. 6e). Particularly notable is that the A-T+ group was reduced from n=37 to n=10 when using a reference protein. Among the ten CSF A-T+, most were close to the decision boundary of being A-T-and only one was classified as A-T+ by PET, indicating that this group could be even further reduced.

**FIGURE 6:**
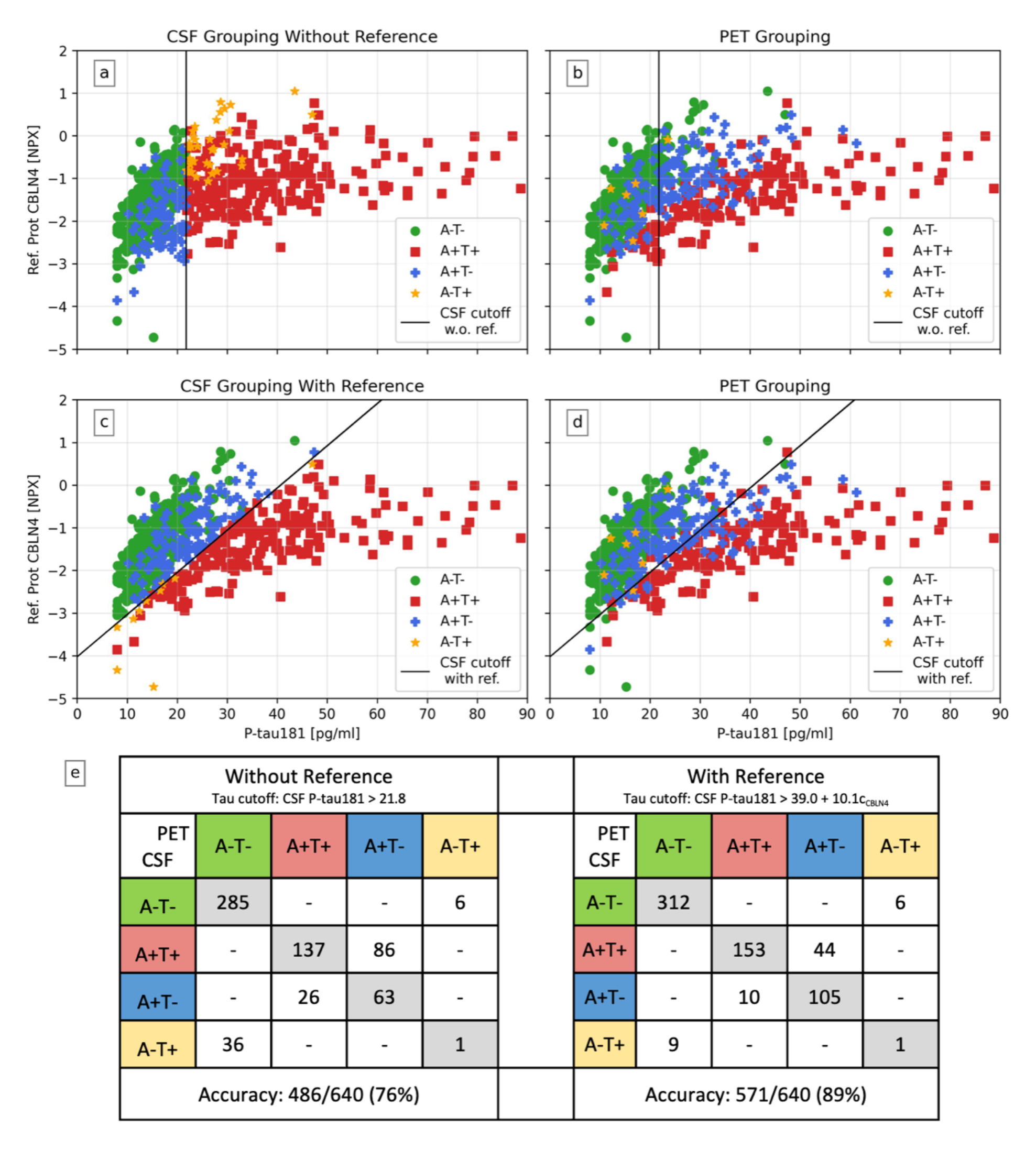
Adjusting for an individual reference protein (here CBLN4) results in a better AT(N) grouping concordance between CSF P-tau181 and tau-PET. In **a)**-**d)**, the x-axis is a participant’s CSF P-tau181 concentrations, and the y-axis the suggested reference protein CBLN4. In **a)** and **c)**, CSF P-tau181 and CSF Aβ42/Aβ40 (cutoff 0.08) has been used to group participants. In **b)** and **d)**, a tau-PET composite corresponding to Braak I-IV with ROIs > 1.36 and CSF Aβ42/Aβ40 was used to group participants. To create a cutoff for CSF P-tau181 (black line), the reference protein CBLN4 was adjusted for in **c)** and **d)** (cutoff: *CSF P-tau181 > 39.0 + 10.1c_CBLN4_*) but not in **a)** and **b)** (cutoff: *CSF P-tau181 > 21.8*). The concordance between CSF P-tau181 and tau-PET grouping increased when not requiring a vertical cutoff line but allowing for it to have a slope. In **e)** corresponding concordance matrices of PET and CSF with and without using a reference for CSF P-tau181 can be seen. The concordance increased from 76% to 89% when adjusting for the reference protein. Particularly notable is that the A-T+ group (which is pathophysiologically difficult to explain) was reduced from n=37 to n=10 when using a reference protein, again in higher concordance with grouping with PET.

Examples of how previously published research results of P-tau181^18, 19^ were affected by this CSF AT(N) grouping improvement can be seen in Fig. 7a and *Supplementary Results: Adjusting for a Reference in P-tau181 Applications*. Fig. 7a and Supplementary Fig. 7-9 show how concentrations of sTREM2, sAXL and sTyro3 turned out to be substantially less differentiable between AT(N) groups when adjusting for a reference protein during grouping. These latter results were clearly more similar to results obtained when using PET (instead of CSF) to define AT(N) groups, indicating that the reported relations between AT(N) and these microglia-related proteins were strongly driven by the mean CSF protein level. Further examples of this effect can be seen in Supplementary Tab. 3 and 4, where correlations between P-tau181 and sTREM2, sAXL, sTyro3 and α-synuclein were clearly attenuated when adjusting for a reference protein.

**FIGURE 7:**
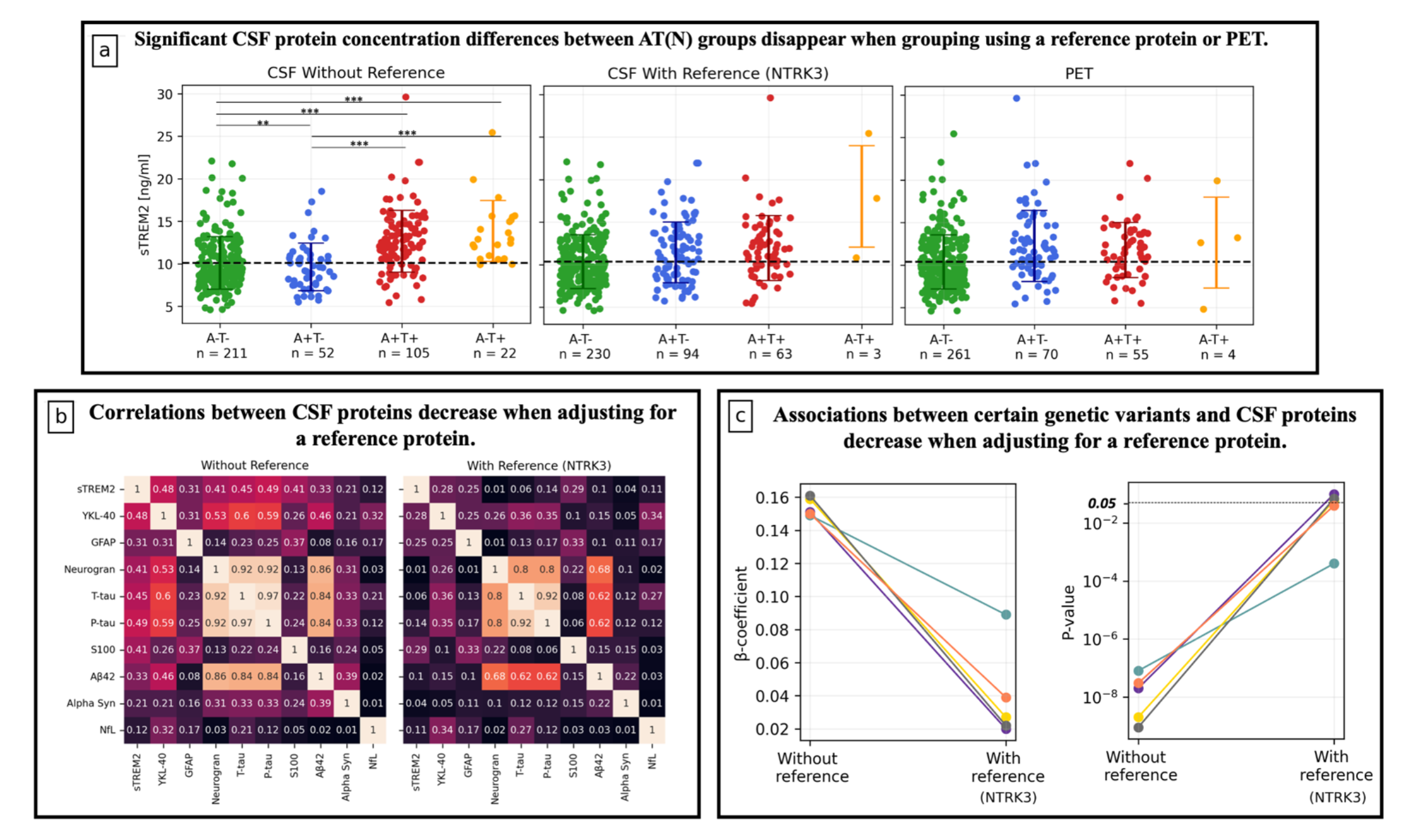
Several CSF proteins are affected by adjusting for a reference protein. Further details of these results and extensive analyses on similar findings can be found in the Supplementary. **a)** During AT(N) grouping assessed by CSF, adjusting for a reference protein created a better concordance with PET when comparing concentration differences of CSF sTREM2. The significant findings from not using a reference protein during grouping most likely appeared due to mean CSF protein level differences between groups. This analysis included NC, SCD and MCI BF2 participants. *P*-values (adjusted for multiple comparisons) were assessed by a one-way ANCOVA adjusted for age and sex. **b)** Partial correlation matrices for ten NeuroToolKit proteins in BF2 cognitively unimpaired Aβ-negative participants with and without adjusting for a reference protein, always adjusting for age and sex. Proteins were sorted according to decreasing association with mean CSF level (see Supplementary Tab. 9). Almost all correlations were severely reduced when adjusting for a reference protein, most clearly seen for proteins highly associated with the mean CSF level (top rows). **c)** Results from protein quantitative trait loci (pQTL) analyses where associations between certain CSF proteins and genetic variants have been identified. Several CSF trans-pQTL associations of the GMNC-OSTN region showed severely weakened relationships when adjusting for a reference protein. For details, see *Supplementary Results: CSF pQTL Analysis* and Supplementary Tab. 7. These models included BF1 participants (n=1445) and were adjusted for age, sex, dementia diagnosis and ten genetic principal components.

### Reference proteins often attenuate CSF biomarker associations

Examples of how other CSF proteins are affected by adjusting for a reference protein are seen in Fig. 7b-7c and *Supplementary Results: Change in Results when Adjusting for Reference Proteins*. In Fig 7b and Supplementary Fig. 10, partial correlations between ten established biomarkers from the NeuroToolKit assay panel proteins with and without adjusting for a reference are given. In general, correlations decreased when adjusting for a reference. This was seen most evidently for the cognitively unimpaired Aβ-negative participants and for proteins highly associated with the mean CSF level, such as CSF levels of sTREM2, YKL-40 and tau. Additionally, examples of reference proteins’ impact on associations between certain CSF proteins and genetic variants are presented. This includes strengthened associations of apolipoprotein E (*APOE*) ε4 alleles with protein levels of ApoE4 and reduced/disappeared associations of trans-protein quantitative trait loci (pQTL) with genes from the GMNC-OSTN region (previously shown to be associated with variations in ventricular volume and suggested to be linked to both CSF P-tau and several other CSF proteins^14, 37^), Fig 7c and Supplementary Tab. 5-7.

## Discussion

We establish the existence of individual mean CSF protein levels, which explain a considerable part of variation in CSF biomarkers. We conclude that many proteins are affected by individual mean CSF levels, including proteins relevant in AD, and may benefit from adjusting for this when used as biomarkers. We identify a robust subset of potential reference proteins (“cluster 11”) from which we further characterize six specific reference protein candidates (NTRK3, NTRK2, BLMH, CBLN4, PTPRN2 and PTPRS) that can significantly improve the accuracy of key AD biomarkers. The results are validated on unseen test data and in an independent cohort. We also provide evidence that Aβ40 works well as a reference protein, not only for Aβ42, but for P-tau181 and other CSF biomarkers as well. Further, we show that several previously reported CSF biomarker classifications and associations were greatly diminished when adjusting for reference proteins. This implies that future studies should account for a reference protein to ensure that observed CSF biomarker relationships are not mainly driven by differences in mean CSF protein levels. Our work focuses on biomarkers in AD, but since the issue of mean CSF protein levels is not AD specific, this concept likely has broad relevance across all neurological and psychiatric conditions where CSF biomarkers are used. The existence of individual variation in mean CSF protein levels provides valuable insights on a CSF characteristic that must be acknowledged and further explored by the field of CSF biomarkers. When investigating potential biological mechanisms driving the mean CSF protein level differences, we found that an increased mean CSF protein level was seen in males and was strongly associated with higher age. This may be connected to other sex and age-related CSF dynamics, like differences in CSF pressure and CSF production and clearance rates. These dynamical differences have previously been observed in both human and animal studies.^38–41^ A reduced CSF production and clearance rate, as seen during aging, may for example contribute to longer accumulation time of CSF proteins, resulting in an increased individual mean CSF protein level. Additionally, as the mean CSF protein level was strongly associated with ventricular volume (but not intracranial volume) when adjusting for age and sex, we believe that CSF dilution is an important factor explaining why these individual differences exist. As CSF fills the ventricles, it is reasonable that the size of the ventricles affects the produced CSF volume independently of CSF protein secretion, leading to these CSF dilution differences.

Supporting this hypothesis is that individuals with idiopathic normal pressure hydrocephalus (iNPH), characterized by an abnormal buildup of CSF resulting in enlarged ventricles, also show substantially low (diluted) CSF AD biomarker levels compared to healthy subjects.^42^

The results of our study challenge previously held notions of strong relationships between several CSF biomarkers in AD. We show that many of these correlations were mainly driven by differences in mean CSF protein levels, as they did not remain robust when accounting for a reference protein. Associations were markedly reduced for P-tau181 versus microglia-related proteins (e.g. sTREM2 or TAM receptors [sAXL and sTYRO3]), astrocytic biomarker YKL-40 and the Parkinson’s disease-related biomarker α-synuclein, indicating that these biological processes/pathologies may not be as related to tau-pathology as both we and others have previously suggested.^18, 19, 27–29^

Our findings provide important insight of how biased intercorrelations between CSF biomarkers and biomarker groups can appear when not accounting for non-disease related differences in mean CSF protein levels. Specifically highlighting this was how the use of a reference protein for P-tau181 in the AT(N) grouping context (Fig. 6e) showed that the A-T+ group (which is pathophysiologically difficult to explain^15^) was reduced from n=37 to n=10 when using a reference protein. This result is more in line with studies using PET to classify individuals according to the AT(N) system.^43^ The characteristics of A-T+ is highly researched and discussed^23–26, 44–46^. We show that this group largely consists of individuals with high overall mean CSF protein levels (rather than any specific disease marker), which may explain why many A-T+ individuals have high CSF concentrations of other proteins than P-tau181^18, 19, 25^. Other conclusions from results in the CSF-based A-T+ group, like hypotheses about tauopathy (T+) not affecting cognition ^47^, can be highly influenced by erroneous classification of individuals with high overall CSF levels into an A-T+ group.

Throughout the evaluation, three general reference proteins (NTRK3, NTRK2 and BLMH) and Aβ40 were examined. For all P-tau181 associations, NTRK3, NTRK2 and Aβ40 performed similarly, while BLMH performed slightly inferior. As reference to Aβ42, Aβ40 was superior. Aβ40 consistently provided improved accuracy and top performance when used as reference for P-tau181 as well, congruent with Guo et al.^13^ As Aβ40 is already often measured in CSF studies, the performance of Aβ40 as a reference can easily be further validated for other biomarkers and cohorts. Aβ40 could be a suitable first individual reference to adjust for when working with CSF AD biomarkers.

We hypothesized that each CSF biomarker might have one or several *optimal* reference proteins that co-vary highly with the main predictor during normal physiology but not in disease, in addition to representing an individual mean CSF protein level. This idea is strengthened by the fact that the top reference protein candidates showed high correlation with the main predictors in cognitively unimpaired Aβ-negative participants but were not predictive of the outcome when applied in models alone. Additionally, simply adjusting for the mean CSF level never resulted in best performance. For Aβ42, a reference that will outperform Aβ40 is unlikely to emerge, as this peptide so closely follows the same biological pathway as Aβ42. Previous work has shown the benefit of adjusting Aβ42 levels with Aβ40^31, 32^, and in extension, our results confirm that Aβ40 performed best as reference for Aβ42 when compared to other possible CSF references. For P-tau181, no such optimal reference has previously been suggested, but three possible candidates (CBLN4, PTPRN2 and PTPRS) were evaluated in this paper. The candidates performed best for two P-tau181 models, but these candidates were only evaluated in BF2 and did not significantly outperform other suggested candidates. Additionally, none of them are as obviously associated with P-tau181 in regards to biological pathway as Aβ40 is with Aβ42, which further decreases their probability of being optimal P-tau181 specific references. However, other factors could make CSF protein concentrations co-vary, such as cellular localization as well as protein size, charge, and solubility. The properties and generalizability of these candidates should be further examined before they can be claimed optimal or non-optimal reference proteins for P-tau181.

As seen in *Supplementary Reference Candidates*, the six novel reference protein candidates (NTRK3, NTRK2, BLMH, CBLN4, PTPRN2 and PTPRS) all had higher association with the mean CSF protein level (partial correlations 0.62-0.80) than Aβ40 (partial correlation 0.48). Many of them were cell surface receptors and involved in cell survival and differentiation. Most had enhanced brain specificity, but low regional brain specificity. From their performance as reference proteins in BF2, we cannot conclude that these specific six candidates were significantly superior reference proteins. On the contrary, several proteins shared similar expression characteristics that were beneficial for a reference protein. We therefore presented a subset of many such proteins by using a data-driven approach to group them into the cluster referred to as “cluster 11”. From the cell type expression analysis, we found that cluster 11 was representative of the entire set of 2,944 proteins with majority of proteins highly expressed in neuronal cells. The proteins of cluster 11 were enriched on cell surfaces and membranes and are therefore probably constantly shedded into the CSF during normal physiology, which can explain why they maintain a relatively consistent concentration level representative of the mean CSF protein level.

A potential limitation of the study was that we only had the possibility to validate parts of the results in an independent cohort. While being a key asset for the data driven approach in the BioFINDER-2 study, the extensive CSF measurements of 2,944 proteins also limited the validation possibilities of the findings. Other cohorts with such extensive CSF measurements are difficult to reproduce and access due to financial and technical constraints. Additionally, the biological and technical variability of the suggested reference protein candidates should be further examined to ensure robustness of longitudinal measurements in participants. Nevertheless, this work supports the proof-of-principle that adjustment for inter-individual differences in mean CSF protein levels will likely be highly useful in future studies aiming to understand associations between different CSF proteins or using key CSF proteins as diagnostic or prognostic biomarkers.

## Conclusion

We show that inter-individual differences in mean CSF protein levels confounds diagnostic and prognostic performance for several CSF biomarkers. These differences can also result in false conclusions regarding associations between different CSF proteins or their relations to genetic variations. The issue can be addressed by using certain CSF reference proteins to represent the non-disease related concentration of the protein studied. Aβ40 is one of several promising general reference proteins (not just for Aβ42) and may be a suitable reference option due to its frequent availability in AD cohorts. Accounting for a CSF reference protein in future studies may help ensure that reported correlations between CSF proteins are not mainly due to mean CSF protein level differences. Our novel reference protein method improves the accuracy of CSF biomarkers, and reduces the risk for false positive findings, with broad implications for both research and clinical practice.

## Methods

### Participants

Two study cohorts were included: the Swedish BioFINDER-2 (BF2) cohort (enrollment from 2017 and still enrolling, n=982, NCT03174938) and the Swedish BioFINDER-1 (BF1) cohort (enrollment between 2010 and 2015, n=1571, NCT01208675). All participants were recruited at Skåne University Hospital and the Hospital of Ängelholm, Sweden. BF2 and BF1 consisted of individuals with either normal cognition (NC), subjective cognitive decline (SCD), mild cognitive impairment (MCI), dementia or another neurodegenerative disease. Conversion to AD dementia was determined during follow-up based on the treating physician’s assessments.^48^ Participants labeled as “non-converted” remained NC, SCD or MCI stable for at least two years. Further details about BF2 and BF1 can be found in ^49^ and ^50^ respectively, or at www.biofinder.se.

Only BF2 participants with complete CSF measures of 2,944 proteins were included for analyses. No other exclusion criteria were implemented, but further filtering was later performed depending on the variables included in the statistical model. In BF1, the participants with complete CSF measures of 369 proteins were included in a similar manner.

### Ethics

The studies were approved by the Swedish Ethical Review Authority, and all participants gave written informed consent to participate.

### CSF Collection and Analysis

CSF samples were collected close in time after baseline examination (first visit) and handled according to established preanalytical protocols.^49, 51^ CSF samples were analyzed with validated, highly sensitive and specific Proximity Extension Assay (PEA) developed by OLINK Proteomics (Uppsala, Sweden). For BF2, the full OLINK Explore 3072 library was used, resulting in eight Proseek Multiplex panels (Oncology I and II, Neurology I and II, Cardiometabolic I and II, Inflammation I and II) to measure the concentration of 2,943 CSF proteins. Each panel contained 367-369 proteins. For BF1, four panels (Neurology-exploratory, Neurology-I, Inflammation-I and Cardiovascular-III) were used to measure the concentration of 368 CSF proteins. Each panel contained 92 proteins. All 368 proteins from the BF1 panels were also included in the BF2 Explore 3072 panels. Protein concentrations were provided as log_2_ scale of Normalized Protein eXpression (NPX) values. High NPX values correspond to high protein concentrations. Details of OLINK quality control and protein detectability are included in *Supplementary Methods*.

CSF biomarkers from the NeuroToolKit assay panel (P-tau181, Aβ42, Aβ40, sTREM2, YKL-40, GFAP, neurogranin, T-tau, S100, alpha synuclein and NfL) were measured in both cohorts using Elecsys assays in accordance with the manufacturer’s instructions (Roche Diagnostics International Ltd).^52^ CSF analyses were performed by technicians blinded to all clinical and imaging data. CSF amyloid positivity was defined based on CSF Aβ42/Aβ40 that was dichotomized using the previously established cutoff of < 0.08.^36^

### PET imaging

In both BF1 and BF2, amyloid-PET imaging was performed using [^18^F]flutemetamol. Standardized uptake value ratio (SUVR) images were created for the 90–110 min post-injection interval with whole cerebellum as reference region. A global neocortical composite region (volume of interest) corresponding to a set of cortical regions was used to summarize [^18^F]flutemetamol data, as described in ^43^. SUVR values were transformed into centiloids. The composite was used as a dichotomous variable with centiloids >20 regarded as amyloid positivity.^53^ In BF2, tau-PET was correspondingly performed using [^18^F]RO948. SUVR images were created for the 70-90 min post-injection interval using the inferior cerebellar cortex as reference region. A composite corresponding to Braak I-IV regions^54^ was used to represent AD-related tau pathology in the brain. The composite was used as a dichotomous variable with SUVR > 1.36 regarded as tau positivity.^55^

### Statistical Analysis

All analyses were implemented using Python version 3.9 or R version 4.2. When searching for reference proteins, all exploratory evaluations were performed using 10-fold-cross-validation within the BF2 training dataset. When evaluating results on the BF2 test dataset, the models were first refit on the full training dataset and thereafter evaluated once on the test dataset. When evaluating results on the BF1 dataset or a new model on the BF2 training dataset, bootstrap-resampling with replacement (n_iter_ = 2,000) was performed such that a resampled training set of same size as the full dataset was created. Thereafter, a validation dataset was created from the participants that were never selected into the training set, which consequently varied in size between runs. This methodology was used to gain higher diversity between runs so that uncertainty estimations within a model could be performed with high reliability.

All protein concentrations were z-scored to allow for comparisons between measures in different units. Standardization was performed within each cohort and always fitted to training data. For every participant, a mean CSF level was computed as the average z-score over all 2,943 OLINK proteins + Aβ40 for BF2, and all 368 OLINK proteins + Aβ40 for BF1. For data exploration and visualization purposes, t-distributed stochastic neighbor embedding (t-SNE)^34^ was applied. t-SNE is formulated as a non-linear optimization problem, aimed to preserve relative similarity of pairwise points in a high dimensional space when projected to a lower one.^34^ The t-SNE results were combined with a semi-supervised K-means clustering^35^ algorithm (K=20) to sufficiently create subsets of data. The supervised part was performed by adjusting K and the random initialization seed so that clear structural clusters were separated and areas with characteristics relevant to a reference protein were joined. This was not a unique nor mathematically optimized way of dividing the t-SNE space. It was solely used due to its efficiency in this application as it enabled a more robust examination of subsets of similarly expressed proteins. To provide evidence that the results were not heavily dependent on the selection of K or random initialization, a robustness analysis is provided, see *Supplementary Methods: K-means Robustness Analysis* and Supplementary Fig. 11-13. Additionally, a sensitivity analysis when only using proteins with missing frequency <75% can be seen in *Supplementary Methods: LOD Sensitivity Analysis* and Supplementary Fig. 14-17, resulting in similar findings as when using all 2,944 proteins.

To analyze associations with a continuous dependent variable, linear regression models were applied. In addition, partial correlation coefficients (Pearson) were computed to study correlation matrices between continuous variables. To predict a dichotomous variable, logistic regression models were applied. As dichotomous data for most models was unbalanced, receiver operating characteristic (ROC) curve and Area under the ROC curve (AUC) were used to evaluate performance. AUCs were compared with a one-tailed ROC test using bootstrapping (n_iter_ = 2,000). To compare AT(N)-groups, one-way ANCOVA was applied. P-values were adjusted for multiple comparisons by Benjamini–Hochberg method. All models were adjusted for age and sex.

### Main models

To search for appropriate reference proteins, performance was evaluated in three logistic regression models. In each model, a well-established AD CSF biomarker (P-tau181 or Aβ42) was used as main predictor of either PET images or conversion to AD dementia. The three models were:

1. **P-tau181→TauPET.** Predicting tau-PET positivity with CSF P-tau181 as main predictor.
2. **Aβ42→AβPET.** Predicting Aβ-PET positivity with CSF Aβ42 as main predictor.
3. **P-tau181→ADDconv.** Predicting conversion to AD dementia versus remained stable for at least 2 years with CSF P-tau181 as main predictor.

The first two models were used to search for suitable reference proteins, the third was solely used for validation. The change in overall model performance when adjusting for different individual references was the measure of interest. A flowchart describing the pre-processing steps after split into training/test data for all models can be seen in Supplementary Fig. 18.

### Cell type expression and pathway enrichment analyses

To investigate properties of a cluster of proteins, we performed pathway enrichment and cell type expression analyses. For pathway enrichment, we used the WEB-based Gene SeT AnaLysis Toolkit (WebGestalt).^56^ We performed a human over-representation analysis (ORA) on cellular components, defining the background set as the 2943 OLINK proteins. For cell type expression, we used Seurat version 4.3.0 to analyze the open-access Human MTG 10x SEA-AD Allen Brain data from 2022.^57^ This dataset includes single-nucleus transcriptomes from 166,868 total nuclei derived from the middle temporal gyrus (MTG) from five post-mortem human brain specimens. We used the class and subclass annotation available from the Allen Institute and applied the function AverageExpression (after removing all “None” annotations). From the average expression we then calculated a percentage expression across all cell types. A bootstrap enrichment test (n=10,000) was used to compare significant (Benjamini–Hochberg corrected q-values <0.05) cell type expression differences between a subset of proteins and all other proteins.

### Data Availability

Anonymized data can be shared to qualified academic researchers after request for the purpose of replicating procedures and results presented in the study. Data transfer must be performed in agreement with EU legislation regarding general data protection regulation and decisions by the Ethical Review Board of Sweden and Region Skåne.

### Code Availability

Code for the analyses can be found in the following GIT repository: https://github.com/karlssonlinda/reference_protein_project. Python dependencies include NumPy^58^, pandas^59^, Matplotlib^60^, Scikit-learn^61^, Statsmodels^62^ and Pingouin^63^. R dependencies include Tidyverse^64^ and pROC^65^.

## Supporting information

Supplementary Results

Supplementary Reference Protein Candidates

## References

1. Nichols, E. et al. Estimation of the global prevalence of dementia in 2019 and forecasted prevalence in 2050: an analysis for the Global Burden of Disease Study 2019. Lancet Public Health 7, e105–e125 (2022).

2. Hansson, O. Biomarkers for neurodegenerative diseases. Nat Med 27, 954–963 (2021).

3. Zetterberg, H. & Blennow, K. Moving fluid biomarkers for Alzheimer’s disease from research tools to routine clinical diagnostics. Mol Neurodegener 16, (2021).

4. Scheltens, P. et al. Alzheimer’s disease. The Lancet 388, 505–517 (2016).

5. van Dyck, C. H. et al. Lecanemab in Early Alzheimer’s Disease. New England Journal of Medicine 388, (2023).

6. Mintun, M. A. et al. Donanemab in Early Alzheimer’s Disease. New England Journal of Medicine 384, 1691–1704 (2021).

7. Mallinckrodt, C. et al. Investigating Partially Discordant Results in Phase 3 Studies of Aducanumab. Journal of Prevention of Alzheimer’s Disease (2023) doi:10.14283/jpad.2023.6.

8. Sakka, L., Coll, G. & Chazal, J. Anatomy and physiology of cerebrospinal fluid. Eur Ann Otorhinolaryngol Head Neck Dis 128, 309–316 (2011).

9. Spector, R., Robert Snodgrass, S. & Johanson, C. E. A balanced view of the cerebrospinal fluid composition and functions: Focus on adult humans. Exp Neurol 273, 57–68 (2015).

10. Johanson, C. E. et al. Multiplicity of cerebrospinal fluid functions: New challenges in health and disease. Cerebrospinal Fluid Res 5, (2008).

11. Tumani, H., Huss, A. & Bachhuber, F. The cerebrospinal fluid and barriers – anatomic and physiologic considerations. in Handbook of Clinical Neurology vol. 146 3–20 (Elsevier B.V., 2017).

12. Bouwman, F. H. et al. Clinical application of CSF biomarkers for Alzheimer’s disease: From rationale to ratios. *Alzheimer’s & Dementia: Diagnosis*, Assessment & Disease Monitoring 14, (2022).

13. Guo, T. et al. Normalization of CSF pTau measurement by Aβ40improves its performance as a biomarker of Alzheimer’s disease. Alzheimers Res Ther 12, (2020).

14. Hansson, O. et al. The genetic regulation of protein expression in cerebrospinal fluid. EMBO Mol Med 15, (2023).

15. Jack, C. R. et al. NIA-AA Research Framework: Toward a biological definition of Alzheimer’s disease. Alzheimer’s and Dementia 14, 535–562 (2018).

16. Bäckström, D. et al. NfL as a biomarker for neurodegeneration and survival in Parkinson disease. Neurology 95, E827–E838 (2020).

17. Düzel, E. et al. Amyloid pathology but not APOE ɛ4 status is permissive for tau-related hippocampal dysfunction. Brain 145, 1473–1485 (2022).

18. Suárez-Calvet, M. et al. Early increase of CSF sTREM2 in Alzheimer’s disease is associated with tau related-neurodegeneration but not with amyloid-β pathology. Mol Neurodegener 14, (2019).

19. Brosseron, F. et al. Soluble TAM receptors sAXL and sTyro3 predict structural and functional protection in Alzheimer’s disease. Neuron 110, 1009–1022.e4 (2022).

20. Delmotte, K., Schaeverbeke, J., Poesen, K. & Vandenberghe, R. Prognostic value of amyloid/tau/neurodegeneration (ATN) classification based on diagnostic cerebrospinal fluid samples for Alzheimer’s disease. Alzheimers Res Ther 13, (2021).

21. Nordengen, K. et al. Glial activation and inflammation along the Alzheimer’s disease continuum. J Neuroinflammation 16, (2019).

22. Ou, Y. N. et al. FDG-PET as an independent biomarker for Alzheimer’s biological diagnosis: A longitudinal study. Alzheimers Res Ther 11, (2019).

23. Soldan, A. et al. ATN profiles among cognitively normal individuals and longitudinal cognitive outcomes. Neurology 92, E1567–E1579 (2019).

24. Pouclet-Courtemanche, H. et al. Frontotemporal dementia is the leading cause of “true” A−/T+ profiles defined with Aβ 42/40 ratio. *Alzheimer’s and Dementia: Diagnosis*, Assessment and Disease Monitoring 11, 161–169 (2019).

25. Oberstein, T. J. et al. Amyloid-β levels and cognitive trajectories in non-demented pTau181-positive subjects without amyloidopathy. Brain 145, 4032– 4041 (2022).

26. Yoon, B. et al. Abnormal tau in amyloid PET negative individuals. Neurobiol Aging 109, 125–134 (2022).

27. Majbour, N. K. et al. Increased levels of CSF total but not oligomeric or phosphorylated forms of alpha-synuclein in patients diagnosed with probable Alzheimer’s disease. Sci Rep 7, (2017).

28. Janelidze, S. et al. CSF biomarkers of neuroinflammation and cerebrovascular dysfunction in early Alzheimer disease. Neurology 91, e867–e877 (2018).

29. Janelidze, S. et al. Cerebrospinal fluid neurogranin and YKL-40 as biomarkers of Alzheimer’s disease. Ann Clin Transl Neurol 3, 12–20 (2016).

30. Tijms, B. M. et al. Pathophysiological subtypes of Alzheimer’s disease based on cerebrospinal fluid proteomics. Brain 143, 3776–3792 (2020).

31. Janelidze, S. et al. CSF Aβ42/Aβ40 and Aβ42/Aβ38 ratios: Better diagnostic markers of Alzheimer disease. Ann Clin Transl Neurol 3, 154–165 (2016).

32. Hansson, O., Lehmann, S., Otto, M., Zetterberg, H. & Lewczuk, P. Advantages and disadvantages of the use of the CSF Amyloid β (Aβ) 42/40 ratio in the diagnosis of Alzheimer’s Disease. Alzheimers Res Ther 11, 1–15 (2019).

33. Zetterberg, H., Blennow, K. & Hanse, E. Amyloid β and APP as biomarkers for Alzheimer’s disease. Exp Gerontol 45, 23–29 (2010).

34. Van Der Maaten, L. & Hinton, G. Visualizing Data using t-SNE. Journal of Machine Learning Research vol. 9 (2008).

35. Hartigan, J. A. Clustering Algorithms. (Wiley, 1975).

36. Pichet Binette, A., et al. Amyloid-associated increases in soluble tau relate to tau aggregation rates and cognitive decline in early Alzheimer’s disease. Nat Commun 13, 6635 (2022).

37. Jansen, I. E. et al. Genome-wide meta-analysis for Alzheimer’s disease cerebrospinal fluid biomarkers. Acta Neuropathol 144, 821–842 (2022).

38. May, C. et al. Cerebrospinal fluid production is reduced in healthy aging. Neurology 40, 500 (1990).

39. Fleischman, D. et al. Cerebrospinal Fluid Pressure Decreases with Older Age. PLoS One 7, (2012).

40. Liu, G. et al. Direct Measurement of Cerebrospinal Fluid Production in Mice. Cell Rep 33, (2020).

41. Preston, J. E. Ageing choroid plexus-cerebrospinal fluid system. Microsc Res Tech 52, 31–37 (2001).

42. Nakajima, M. et al. Guidelines for management of idiopathic normal pressure hydrocephalus (Third edition): Endorsed by the Japanese society of normal pressure hydrocephalus. Neurol Med Chir (Tokyo) 61, 63–97 (2021).

43. Ossenkoppele, R. et al. Amyloid and tau PET-positive cognitively unimpaired individuals are at high risk for future cognitive decline. Nat Med 28, 2381–2387 (2022).

44. Tan, M. S. et al. Longitudinal trajectories of Alzheimer’s ATN biomarkers in elderly persons without dementia. Alzheimers Res Ther 12, (2020).

45. Allegri, R. F. et al. Prognostic value of ATN Alzheimer biomarkers: 60-month follow-up results from the Argentine Alzheimer’s Disease Neuroimaging Initiative. *Alzheimer’s and Dementia: Diagnosis*, Assessment and Disease Monitoring 12, (2020).

46. Delvenne, A. et al. Cerebrospinal fluid proteomic profiling of individuals with mild cognitive impairment and suspected non-Alzheimer’s disease pathophysiology. Alzheimer’s and Dementia (2022) doi:10.1002/alz.12713.

47. Düzel, E. et al. Amyloid pathology but not APOE ɛ4 status is permissive for tau-related hippocampal dysfunction. Brain 145, 1473–1485 (2022).

48. Palmqvist, S. et al. Prediction of future Alzheimer’s disease dementia using plasma phospho-tau combined with other accessible measures. Nat Med (2021) doi:10.1038/s41591-021-01348-z.

49. Palmqvist, S. et al. Discriminative Accuracy of Plasma Phospho-tau217 for Alzheimer Disease vs Other Neurodegenerative Disorders. JAMA - Journal of the American Medical Association 324, 772–781 (2020).

50. Palmqvist, S. et al. Cerebrospinal fluid and plasma biomarker trajectories with increasing amyloid deposition in Alzheimer’s disease. EMBO Mol Med 11, (2019).

51. Palmqvist, S. et al. Accuracy of brain amyloid detection in clinical practice using cerebrospinal fluid β-Amyloid 42: A cross-validation study against amyloid positron emission tomography. JAMA Neurol 71, 1282–1289 (2014).

52. Salvadó, G. et al. Optimal combinations of CSF biomarkers for predicting cognitive decline and clinical conversion in cognitively unimpaired participants and mild cognitive impairment patients: A multi-cohort study. Alzheimer’s and Dementia (2023) doi:10.1002/alz.12907.

53. Amadoru, S. et al. Comparison of amyloid PET measured in Centiloid units with neuropathological findings in Alzheimer’s disease. Alzheimers Res Ther 12, (2020).

54. Cho, H. et al. In vivo cortical spreading pattern of tau and amyloid in the Alzheimer disease spectrum. Ann Neurol 80, 247–258 (2016).

55. Leuzy, A. et al. Diagnostic performance of RO948 F 18 tau positron emission tomography in the differentiation of alzheimer disease from other neurodegenerative disorders. JAMA Neurol 77, 955–965 (2020).

56. Liao, Y., Wang, J., Jaehnig, E. J., Shi, Z. & Zhang, B. WebGestalt 2019: gene set analysis toolkit with revamped UIs and APIs. Nucleic Acids Res 47, W199– W205 (2019).

57. Dataset: Allen Institute for Brain Science (2022). Allen Cell Types Database -- Human MTG 10x [dataset]. Available from celltypes.brain-map.org/rnaseq.

58. Harris, C. R. et al. Array programming with NumPy. Nature 585, 357–362 (2020).

59. Mckinney, W. Data Structures for Statistical Computing in Python. Proceedings of the 9th Python in Science Conference (2010).

60. Hunter J. Matplotlib: A 2D graphics environment. Comput Sci Eng 9, (2007).

61. Pedregosa, F., et al. Scikit-learn: Machine Learning in Python. Journal of Machine Learning Research vol. 12 http://scikit-learn.sourceforge.net. (2011).

62. Seabold, S. & Perktold, J. Statsmodels: Econometric and Statistical Modeling with Python. Proceedings of the 9th Python in Science Conference (2010).

63. Vallat, R. Pingouin: statistics in Python. J Open Source Softw 3, 1026 (2018).

64. Wickham, H. et al. Welcome to the Tidyverse. J Open Source Softw 4, 1686 (2019).

65. Robin, X. et al. pROC: An open-source package for R and S+ to analyze and compare ROC curves. BMC Bioinformatics 12, (2011).

