## Supplementary Results for "Cerebrospinal fluid reference proteins increase accuracy and interpretability of biomarkers for brain diseases"

### Changes in Results when Adjusting for Reference Proteins

In this section, examples of CSF relationships that are strengthened or weakened when adjusting for reference proteins are given. This was done by exploring the available CSF protein data in BF1 and BF2, with analyses often partially or fully inspired from findings in previous publications. The analysis was aimed to contribute to further understanding of CSF biomarkers and how reference proteins can affect certain results. In all examples, either Aβ40, NTRK3 or CBLN4 was used as a reference protein.

#### NeuroToolKit Correlations

For this analysis, the relationships of ten established biomarkers measured on an Elecsys^®^ platform (NeuroToolKit assay panel) in BF2 and BF1 were evaluated. This was done by examining the partial correlation (Pearson) adjusting for age, sex, and a potential reference protein. All biomarkers that were evaluated are summarized in Supplementary Tab. 9, where their association with the mean CSF level is given (sorted in decreasing order). Correlation matrices can be seen in Supplementary Fig. 10, where the effect of adjusting for Aβ40 or NTRK3 in the full datasets and for cognitively unimpaired Aβ-negative participants are provided. The correlation matrices show how several correlations were decreased or increased when adjusting for a reference protein. Decreased correlations were most clearly seen for proteins highly associated with the mean CSF level (top rows).

#### *APOE*4 Genotype vs Protein Expression

The number of apolipoprotein E (*APOE*) ε4 alleles is highly associated with the protein expression of ApoE4. Measurements of ApoE4 CSF protein concentrations were performed using high resolution parallel reaction monitoring mass spectrometry in BF1 for participants with one or two ε4 alleles. Details about the methodology can be found in ^2^. Adjusting for a reference protein strengthens the association between the number of ε4 alleles and protein expression of ApoE4, as can be seen in Supplementary Tab. 5.

#### CSF pQTL Analysis (Association of genetic variants and proteins)

In protein quantitative trait loci (pQTL) analyses, associations between certain CSF proteins and genetic variants (Single Nucleotide Polymorphism, SNP) have been identified. Such an analysis has previously been performed on the BF1 cohort, see ^3^. There, it was shown that brain volumetric measures may be a confounder for CSF trans-pQTL associations of the GMNC-OSTN region, hypothesized to be due to dilution effects on proteins. To further investigate the effect of adjusting for a reference protein for such pQTL associations, five CSF protein associations with genes of the GMNC-OSTN region from the mentioned article were selected. The associations were highly significant when not adjusting for a reference. All proteins that were evaluated are summarized in Supplementary Tab. 6, where their association with the mean CSF level is provided.

The proteins were evaluated in linear regression models, with the CSF protein as outcome and SNP as main predictor. In all models, age, sex, dementia diagnosis and ten genetic principal components were adjusted for. The change in association when also adjusting for a reference protein (Aβ40 or NTRK3) was the measure of interest. The changes in β-coefficients, P-values and R-squared were used to compare the effect of genotype between different models. The result can be seen in Supplementary Tab. 7. In general, the trans-pQTL associations of the GMNC-OSTN were severely weakened/disappeared when adjusting for a reference protein. This trend was stronger when adjusting for NTRK3 than Aβ40, emphasizing NTRK3s potential as a potential reference protein that can generalize over many CSF biomarker applications.

#### Adjusting for a Reference Protein in P-tau181 Applications

Several articles have been evaluating associations between P-tau181 and other CSF proteins. In addition, CSF P-tau181 has been used in AT(N) grouping to compare CSF protein levels between groups. As has been shown in this work, the properties of P-tau181 are affected when adjusting for a reference protein. Here, we aim to exemplify a few cases where adjusting for a reference protein affects the result of previous work. The proteins that were evaluated were included in Supplementary Tab. 9 where their association with the mean CSF level is described.

##### sTREM2, sAXL, sTyro3 and YKL-40 association with P-tau181 in NC, SCD and MCI

In several articles, AT(N) grouping has been performed to compare inter-group differences for CSF proteins. Examples are sTREM2^4^, sAXL^5^ and sTyro3^5^. A-grouping has been done using CSF Aβ42/Aβ40, and T-grouping using CSF P-tau181 without adjusting for any reference proteins. It is likely that there will be a bias of elevated levels of proteins in T+ groups and decreased levels of proteins in T- groups if no reference protein is adjusted for. The reference protein effect is visualized by grouping sTREM2 (Supplementary Fig. 7), sAXL (Supplementary Fig. 8) and sTyro3 (Supplementary Fig. 9) into AT(N) groups and compare group differences for the BF2 cohort of participants with NC, SCD or MCI. Here, T-grouping has been performed in five ways: CSF without reference (P-tau cutoff 21.8 pg/ml as in ^4^), CSF with reference protein Aβ40 (logistic regression, cutoff: *CSF P-tau181 > 8.14 + 1.20c_Aβ40_*), CSF with reference protein NTRK3 (logistic regression, cutoff: *CSF P-tau181 > -6.10 + 11.6c_NTRK3_*), CSF with a reference protein CBLN4 (logistic regression, cutoff*: CSF P-tau181 > 40.3 + 9.27c_CBLN4_*) and PET. If using CSF grouping, adjusting for a reference protein reduced the group differences, creating a better concordance between CSF and PET grouping and removing strong correlations that seem to have appeared due to mean CSF protein levels.

In addition, the three proteins’ association with P-tau181 were evaluated in linear regression models (adjusted for age and sex). Here, the protein YKL-40 was also evaluated as it has been found to be highly associated with P-tau181 in ^6^, both for Aβ- and Aβ+ participants. As seen in Supplementary Tab. 3, the effects of the associations were severely reduced or even disappeared when adjusting for a reference protein. This was particularly evident for sTREM2, sAXL and sTyro3, whereas for YKL-40 the relationship was reduced in significance but still relatively strong.

##### α-synuclein in AD Dementia

In previous research, CSF α-synuclein has been found be highly associated with CSF P-tau181 in AD dementia participants, see for example Majbour et al. and Slaets et al.^7,8^ Therefore, a similar analysis as in Supplementary Tab. 3 was performed for CSF α-synuclein in BF2, but here only for participants with AD dementia, see Supplementary Tab. 4. Again, the effect size of the association decreases when adjusting for reference proteins.

### References

1.	Shahapure, K. R. & Nicholas, C. Cluster quality analysis using silhouette score. in *Proceedings - 2020 IEEE 7th International Conference on Data Science and Advanced Analytics, DSAA 2020* 747–748 (Institute of Electrical and Electronics Engineers Inc., 2020). doi:10.1109/DSAA49011.2020.00096.

2.	Minta, K. *et al.* Quantification of total apolipoprotein e and its isoforms in cerebrospinal fluid from patients with neurodegenerative diseases. *Alzheimers Res Ther* **12**, (2020).

3.	Hansson, O. *et al.* The genetic regulation of protein expression in cerebrospinal fluid. *EMBO Mol Med* **15**, (2023).

4.	Suárez-Calvet, M. *et al.* Early increase of CSF sTREM2 in Alzheimer’s disease is associated with tau related-neurodegeneration but not with amyloid-β pathology. *Mol Neurodegener* **14**, (2019).

5.	Brosseron, F. *et al.* Soluble TAM receptors sAXL and sTyro3 predict structural and functional protection in Alzheimer’s disease. *Neuron* **110**, 1009-1022.e4 (2022).

6.	Janelidze, S. *et al.* CSF biomarkers of neuroinflammation and cerebrovascular dysfunction in early Alzheimer disease. *Neurology* **91**, e867–e877 (2018).

7.	Majbour, N. K. *et al.* Increased levels of CSF total but not oligomeric or phosphorylated forms of alpha-synuclein in patients diagnosed with probable Alzheimer’s disease. *Sci Rep* **7**, (2017).

8.	Slaets, S. *et al.* Increased CSF α-synuclein levels in Alzheimer’s disease: Correlation with tau levels. *Alzheimer’s and Dementia* **10**, S290–S298 (2014).
