## Supplementary Figures for "Cerebrospinal fluid reference proteins increase accuracy and interpretability of biomarkers for brain diseases"

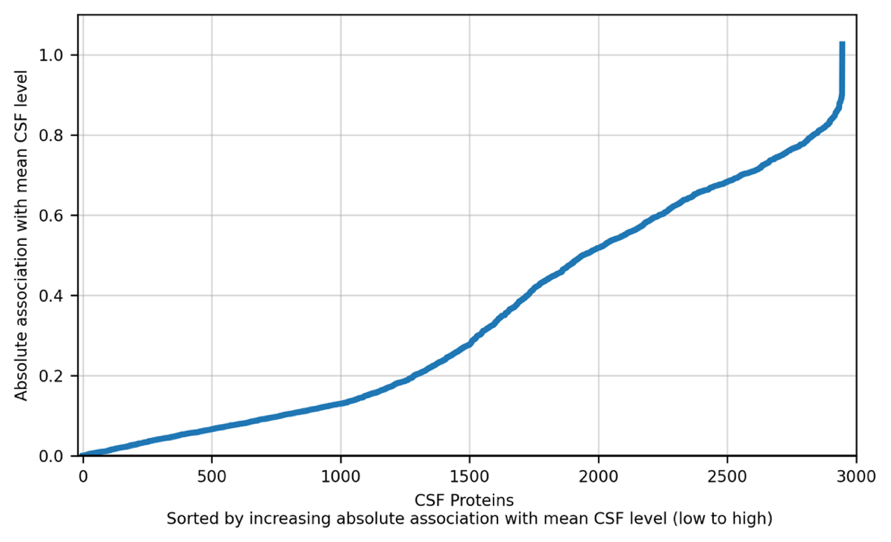

**Supplementary Figure 1**: **Absolute association (β-value) with mean CSF level for all 2944 proteins.** Computed in a linear regression model adjusted for age and sex. The result was used to sort the x-axis in Fig. 2 and Supplementary Fig. 3.

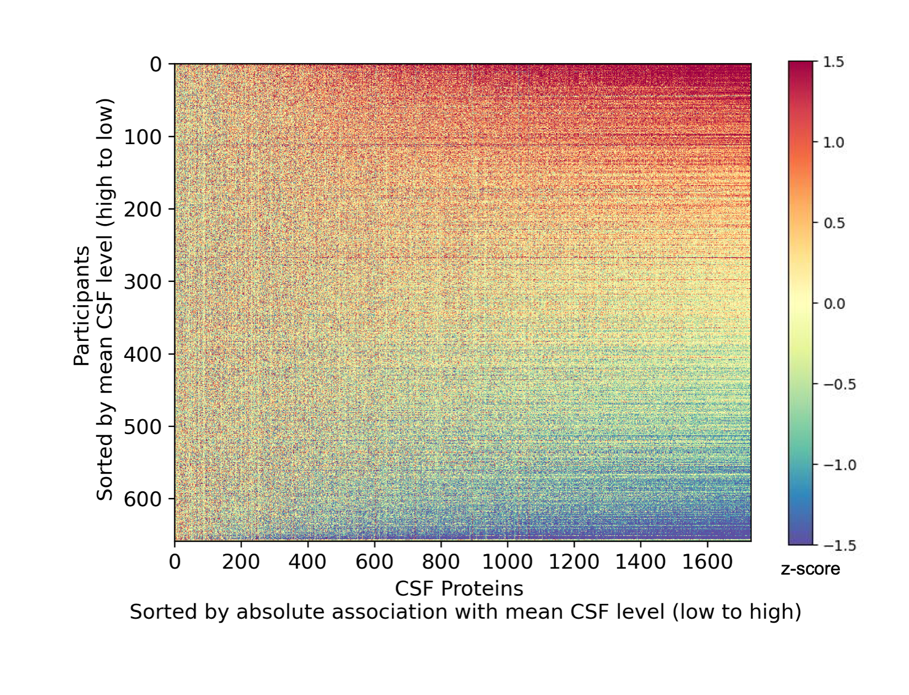

**Supplementary Figure 2: Visualization of individual CSF levels only including proteins with missing frequency < 75%.** For each participant (row), the z-score of 1730 highly detected CSF proteins, sorted by absolute association with mean CSF level, is displayed. As expected, the individual CSF level is prominently shown in highly expressed proteins.

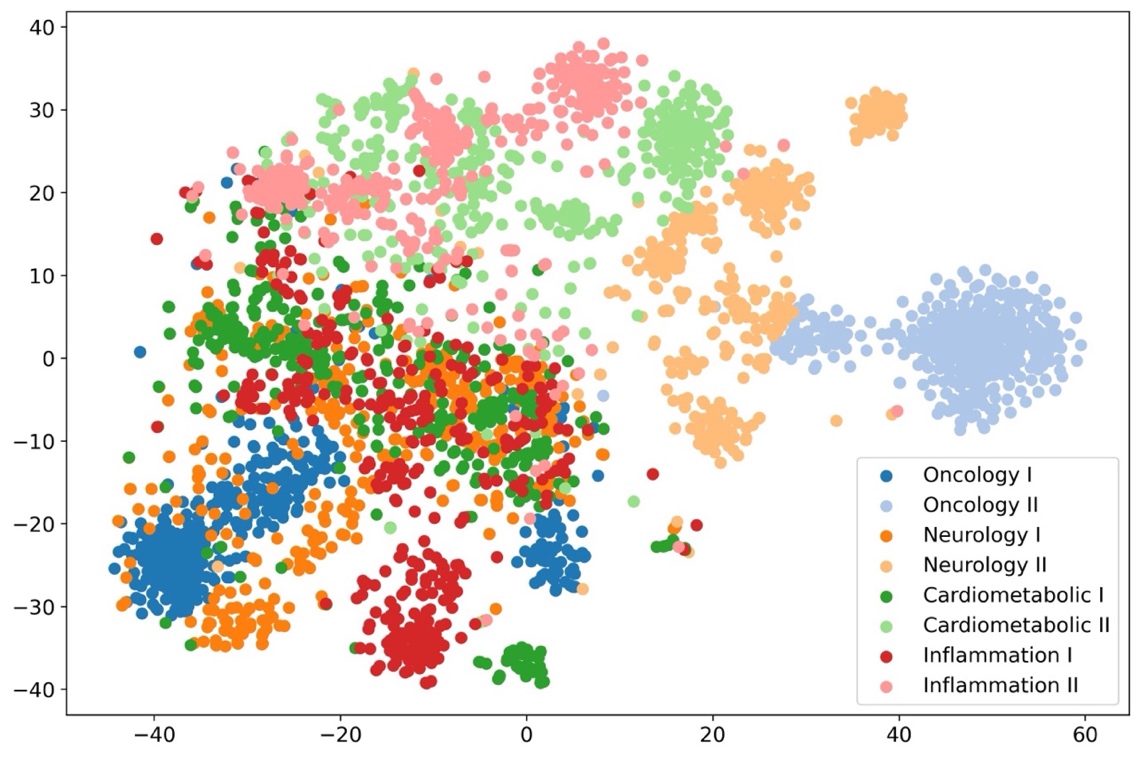

**Supplementary Figure 3**: **t-SNE OLINK panel coloring.** t-SNE dimensionality reduction of 658-dimensional space of participants into a two-dimensional space, colored by OLINK panel inclusion.

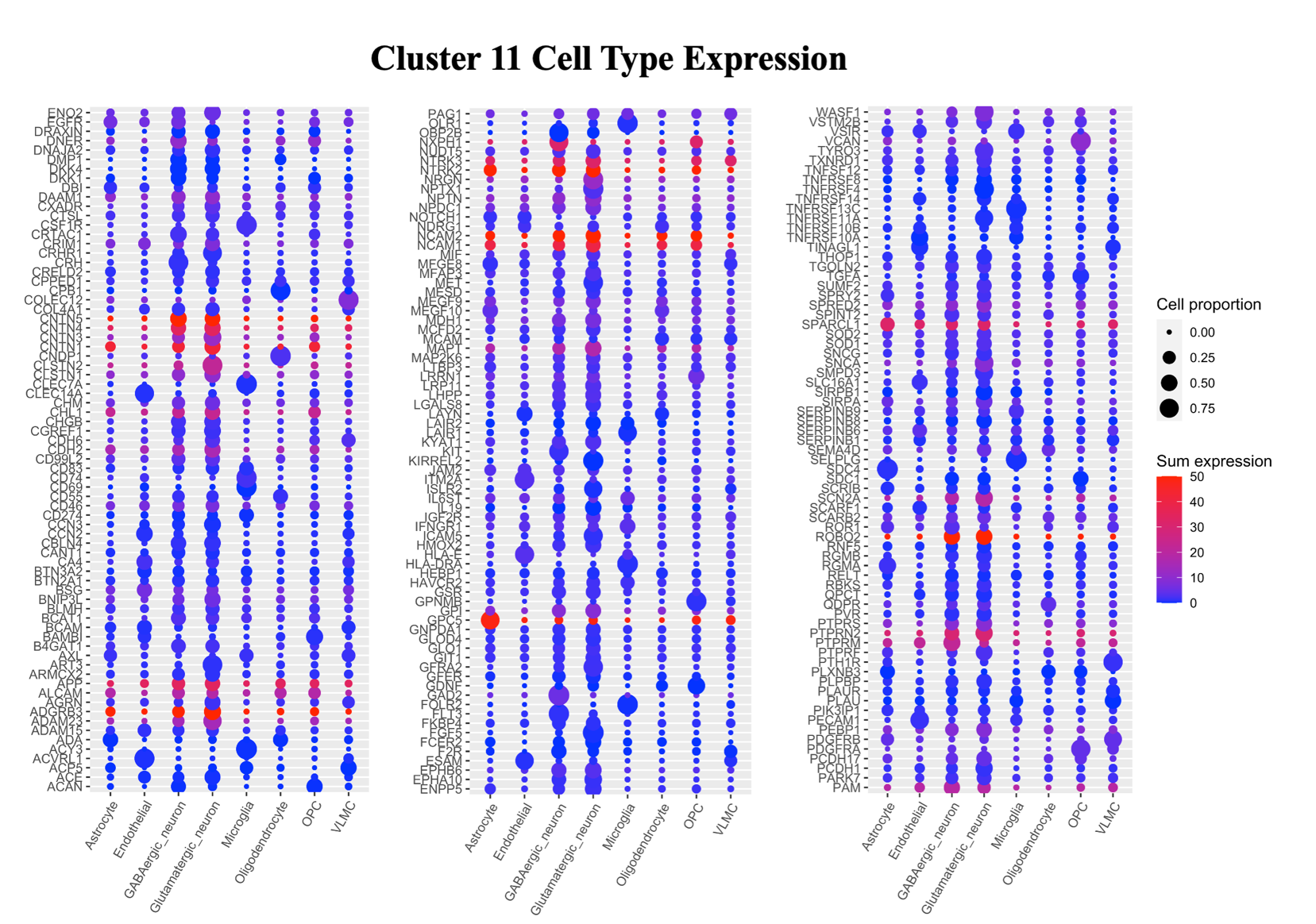

**Supplementary Figure 4**: **Cluster 11 cell type expression.** Average cell type expression profiles for 217 of the 219 genes corresponding to the proteins in cluster 11. Based on data from 5 post-mortem human brain specimens in the Allen Brain Data. Abbreviations: gamma-aminobutyric acid (GABA), Oligodendrocyte progenitor cell (OPC), vascular and leptomeningeal cell (VLMC).

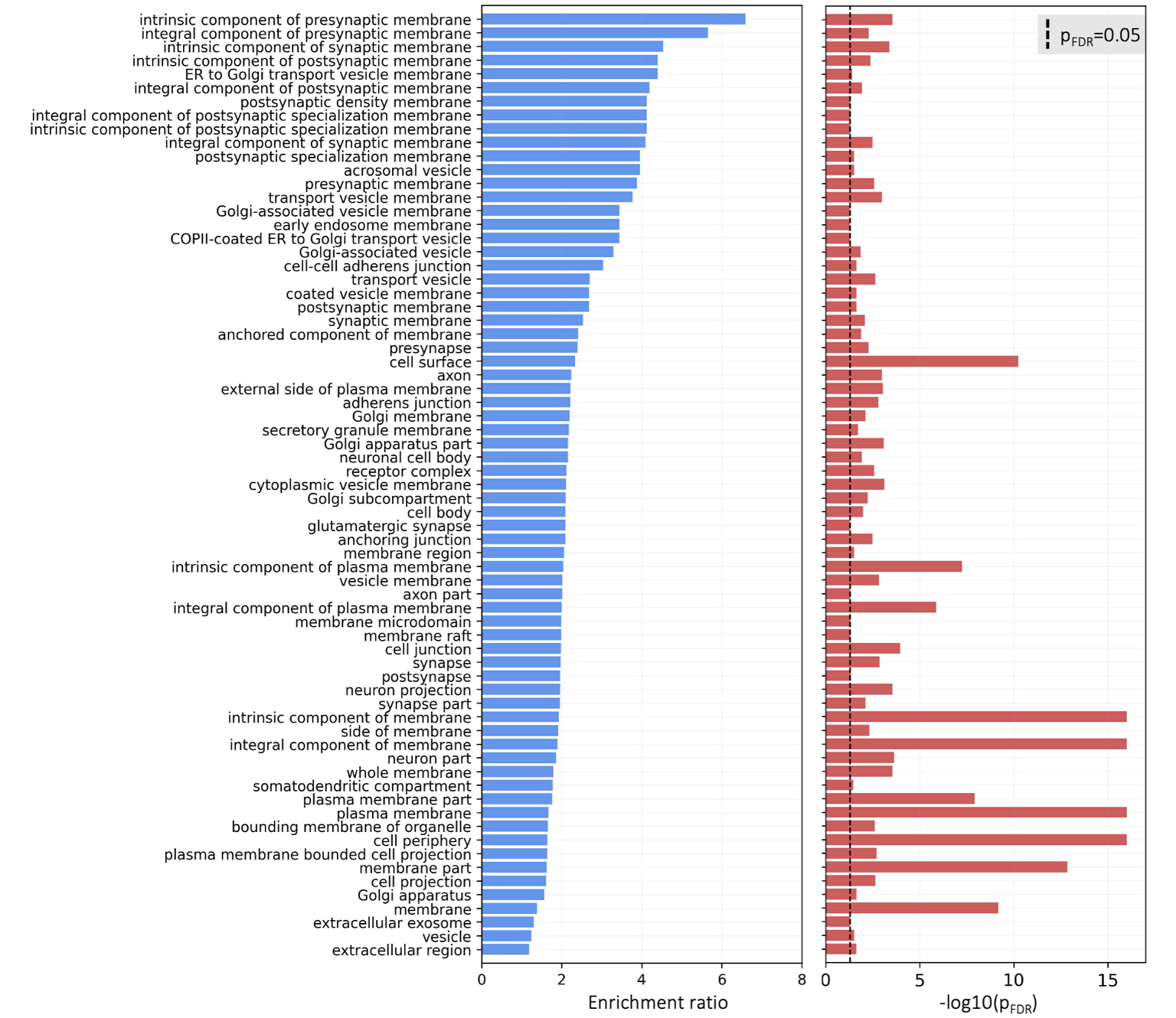

**Supplementary Figure 5**: **Cluster 11 cellular component enrichment analysis.** Over representation cellular component enrichment analysis for 217 of the 219 proteins in cluster 11 performed on the WEB-based Gene SeT AnaLysis Toolkit (WebGestalt). The background set was defined as all 2943 OLINK proteins.

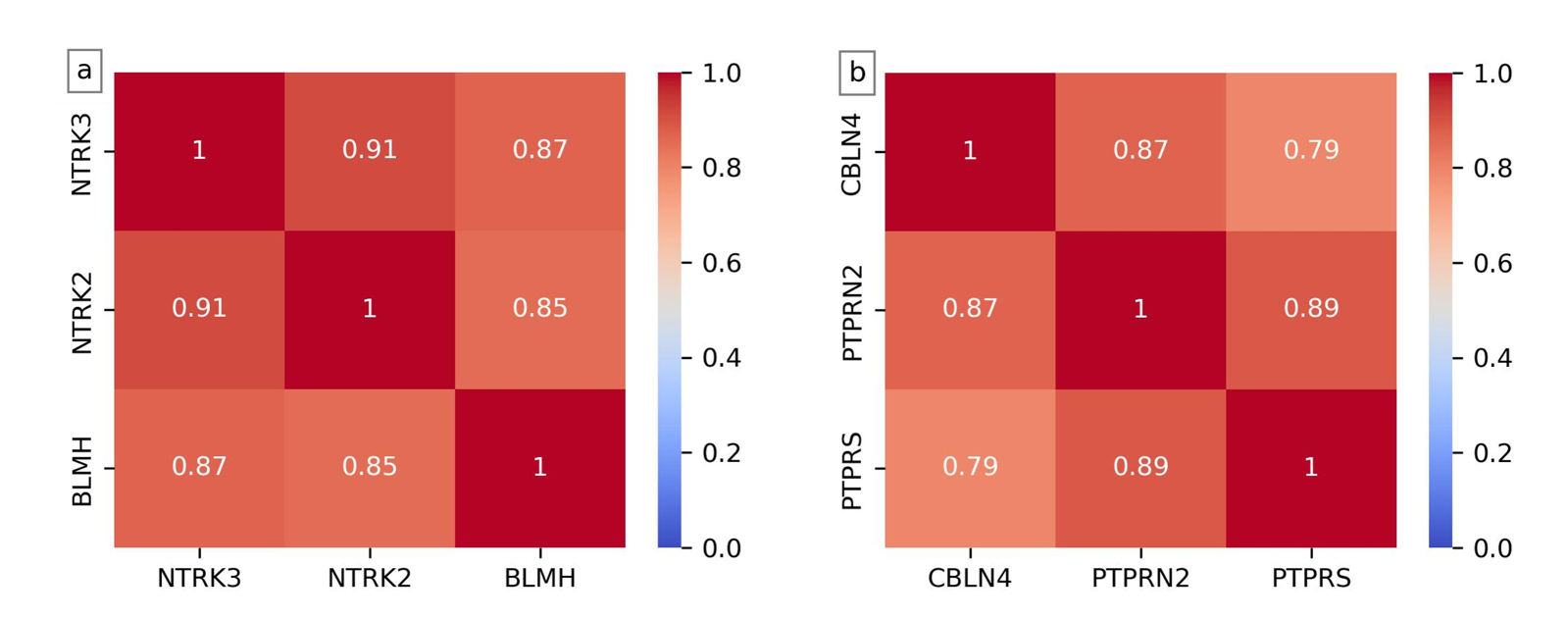

**Supplementary Figure 6: Correlation matrix of general candidates.** Pearson correlation matrix of the **a)** general reference protein candidates NTRK3, NTRK2 and BLMH and **b)** P-tau181 specific reference protein candidates CBLN4, PTPRN2 and PTPRS.

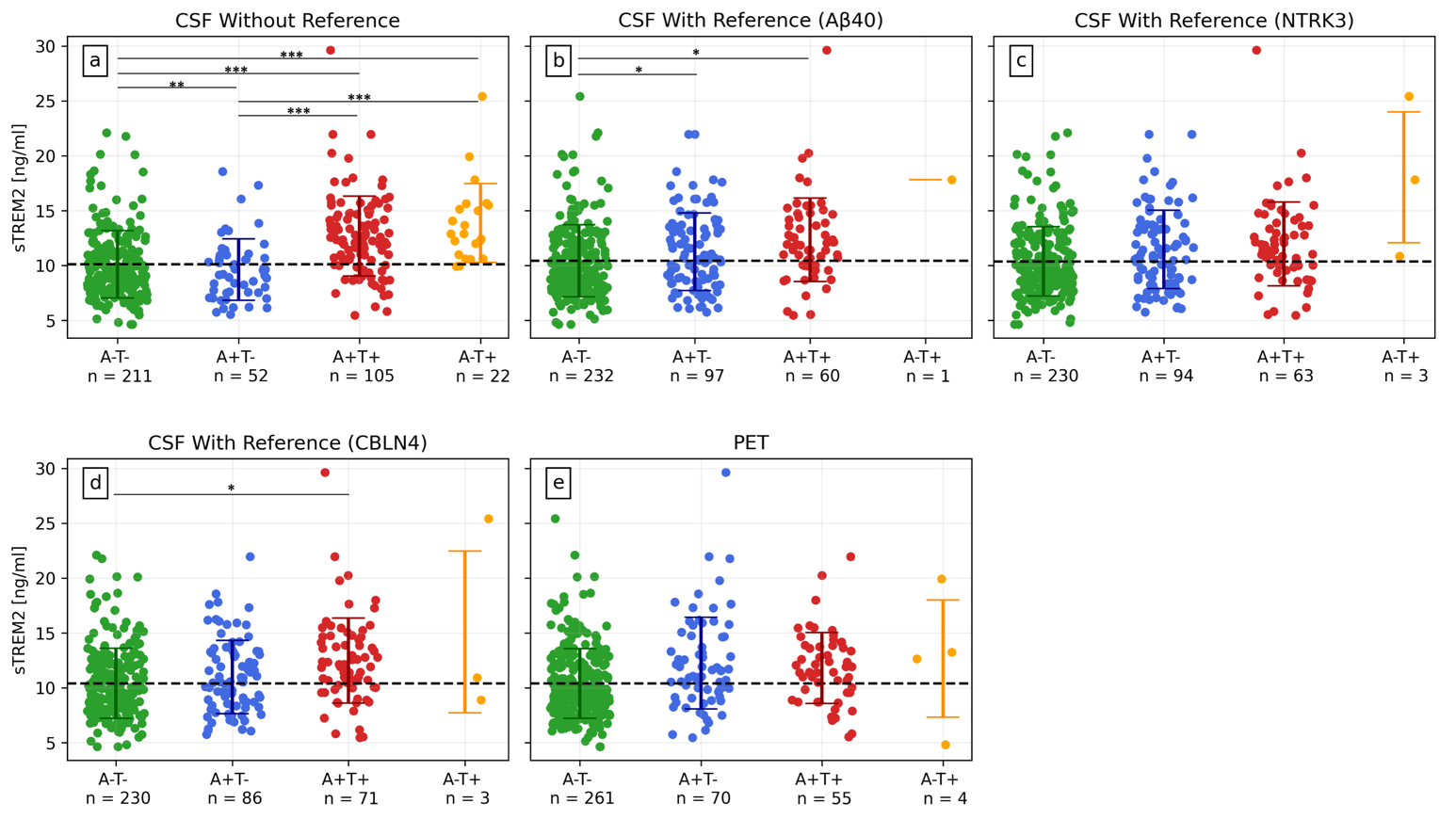

**Supplementary Figure 7: sTREM2 levels by AT(N) grouping for NC, SCD and MCI. a)** CSF grouping without reference. **b)** CSF grouping with Aβ40 as reference. **c)** CSF grouping with NTRK3 as reference. **d)** CSF grouping with CBLN4 as reference. **e)** PET grouping. The differences between groups were affected by how the grouping was performed. If using CSF grouping, adjusting for a reference protein reduced the group differences, creating a better concordance between CSF and PET grouping. *P*-values (adjusted for multiple comparisons) were assessed by a one-way ANCOVA adjusted for age and sex.

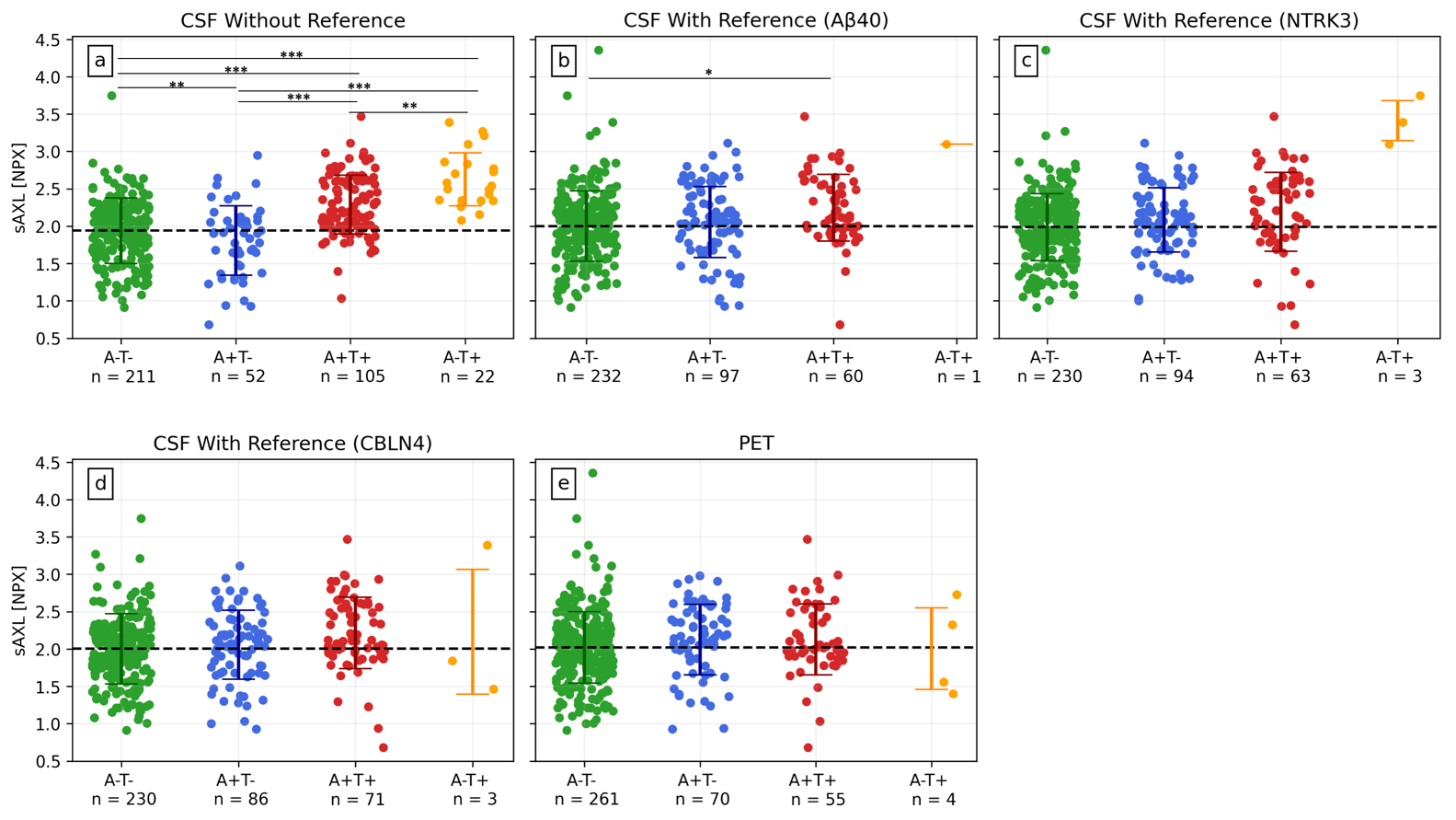

**Supplementary Figure 8: sAXL levels by AT(N) grouping for NC, SCD and MCI. a)** CSF grouping without reference. **b)** CSF grouping with Aβ40 as reference. **c)** CSF grouping with NTRK3 as reference. **d)** CSF grouping with CBLN4 as reference. **e)** PET grouping. The differences between groups were affected by how the grouping was performed. If using CSF grouping, adjusting for a reference protein reduced the group differences, creating a better concordance between CSF and PET grouping. *P*-values (adjusted for multiple comparisons) were assessed by a one-way ANCOVA adjusted for age and sex.

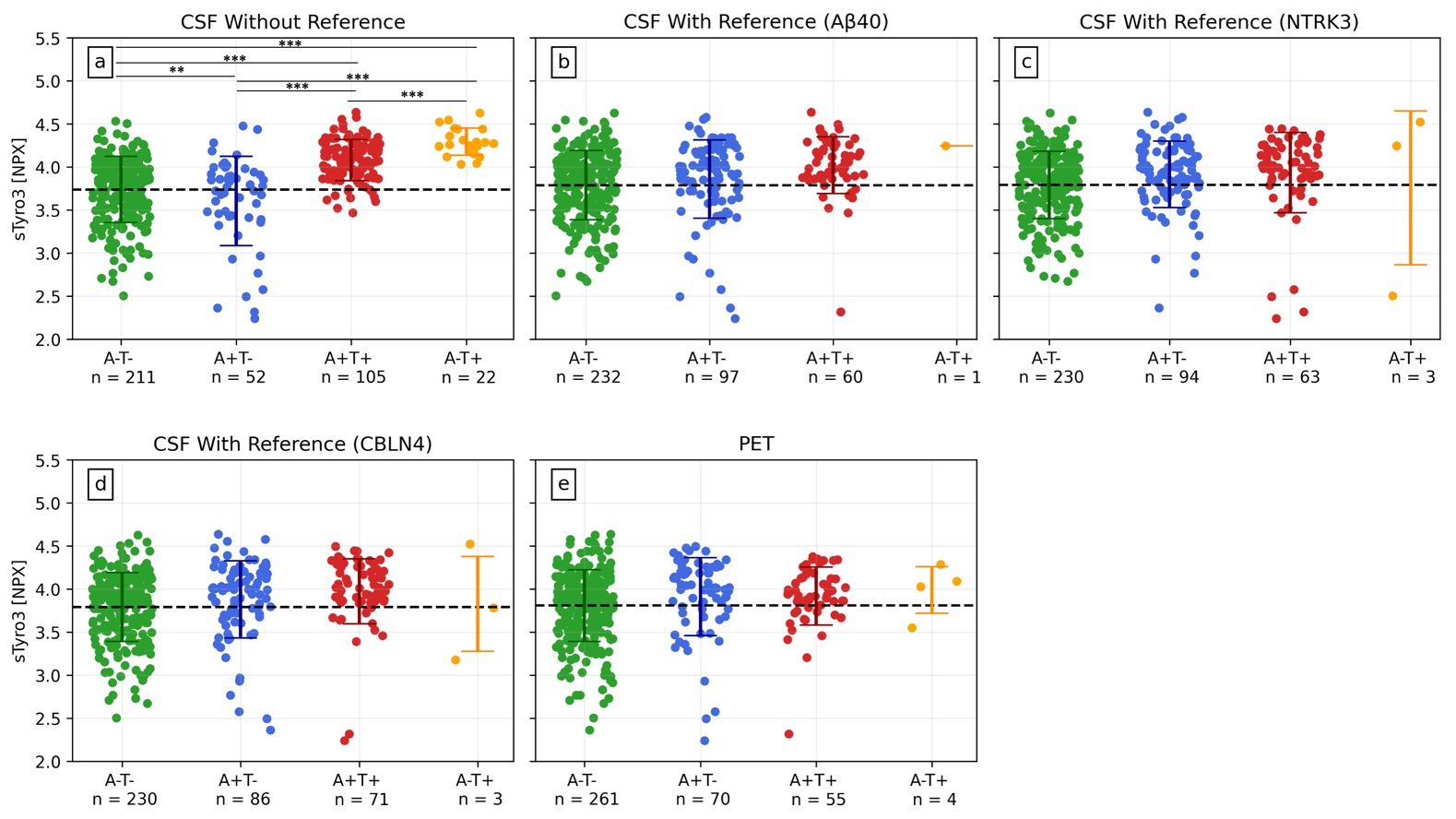

**Supplementary Figure 9: sTyro3 levels by AT(N) grouping for NC, SCD and MCI. a)** CSF grouping without reference. **b)** CSF grouping with Aβ40 as reference. **c)** CSF grouping with NTRK3 as reference. **d)** CSF grouping with CBLN4 as reference. **e)** PET grouping. The differences between groups were affected by how the grouping was performed. If using CSF grouping, adjusting for a reference protein reduced the group differences, creating a better concordance between CSF and PET grouping. *P*-values (adjusted for multiple comparisons) were assessed by a one-way ANCOVA adjusted for age and sex.

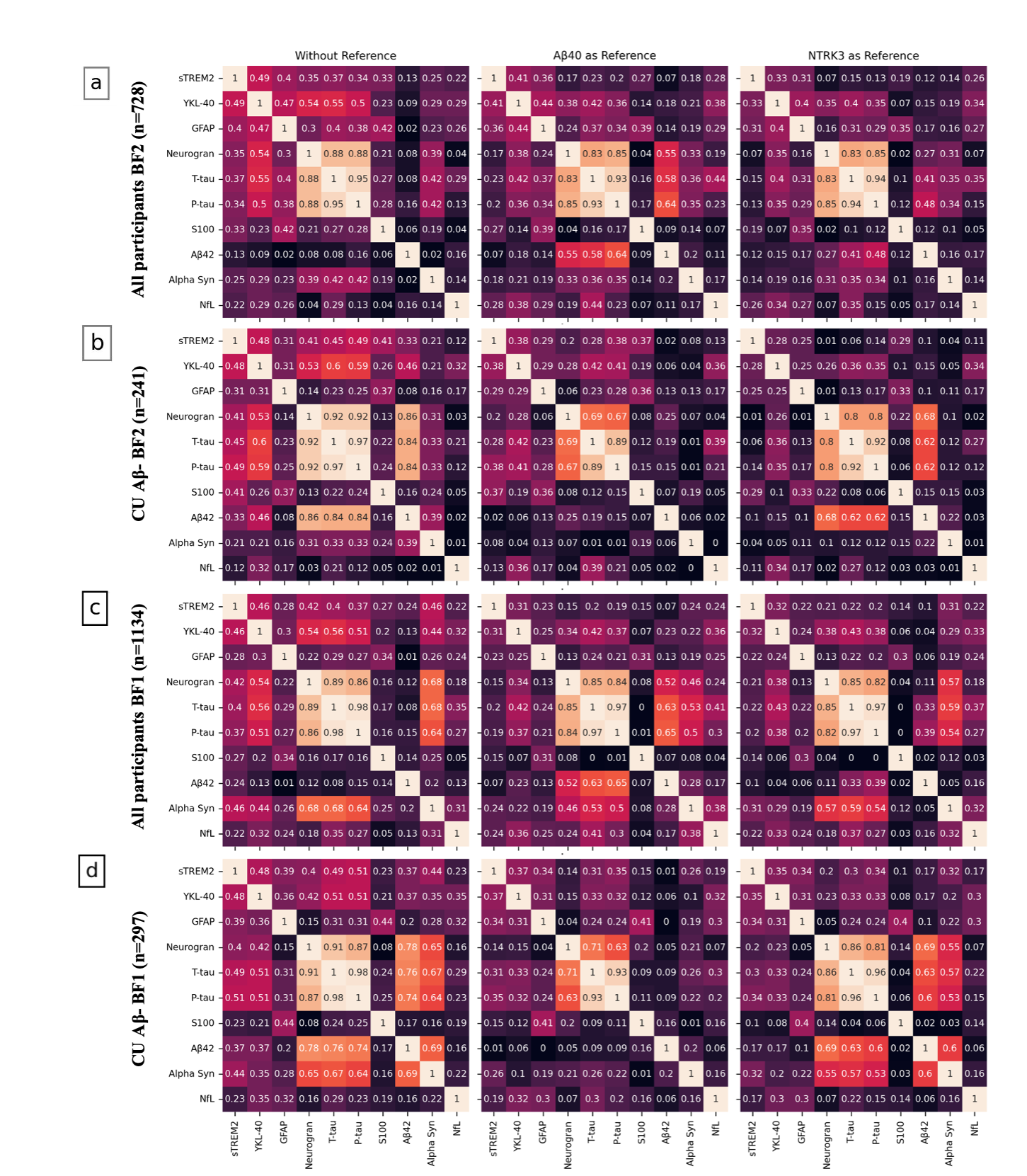

**Supplementary Figure 10: Correlation matrices for ten NeuroToolKit proteins in BF1 and BF2 with and without adjusting for a reference protein.** Partial correlation matrices for ten NeuroToolKit proteins in BF2 training and BF1 datasets, adjusting for age and sex. Proteins were sorted according to decreasing association with mean CSF level (se Supplementary Tab. 9). From these results, a comparison of the change in correlation when also adjusting for a reference protein (Aβ40 or NTRK3) can be made. Almost all correlations were severely reduced when adjusting for a reference protein, particularly evident in cognitively unimpaired individuals without AD pathology. **a)** All participants in BF2 (n=728). **b)** Cognitively unimpaired Aβ-negative BF2 participants (n=241). **c)** All participants in BF1 (n=1134). **d)** Cognitively unimpaired Aβ-negative BF1 participants (n=297).

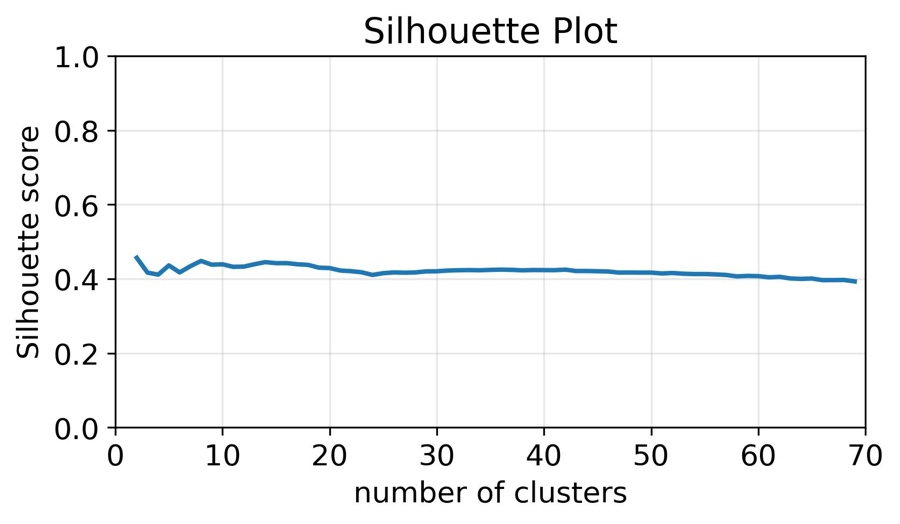

**Supplementary Figure 11: Mean silhouette score for K-means clustering of t-SNE space**. For each K, a score was computed as the average of 20 different random initializations. As seen here, the silhouette score did not vary much between different Ks in the range 2-70, hence not evidently favoring any specific K.

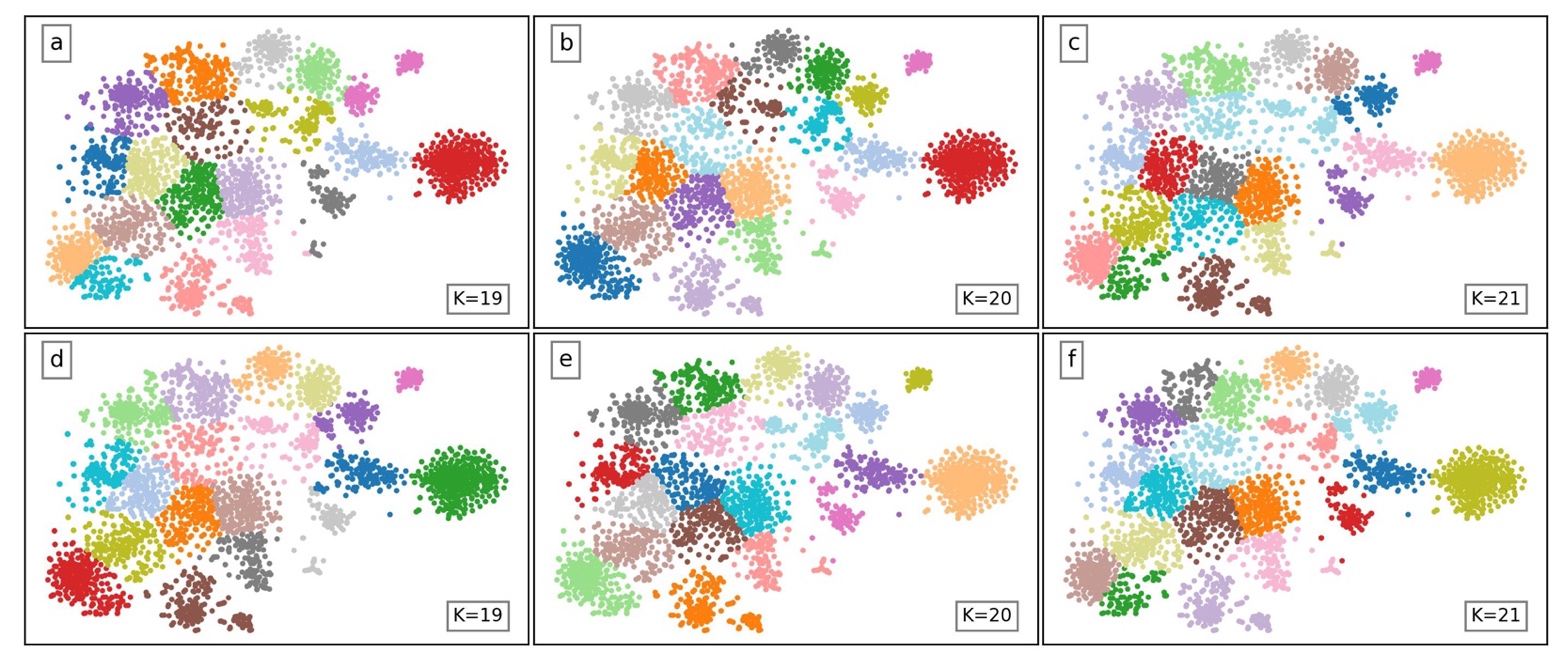
 **Supplementary Figure 12: Semi-supervised K-means clustering of t-SNE space for different Ks and random initialization seeds.** As seen in the six subfigures, some variations in the result are given when changing K and random initialization seeds. The semi-supervised K-means clustering with K=20 is a non-unique nor mathematically optimized tool to select a subset of proteins. Still, it fulfills the purpose of adding robustness when analyzing expression characteristics rather than single proteins in a reproducible way. This could also have been achieved with different Ks and random initialization seeds as seen in the figure, making the results highly reproducible.

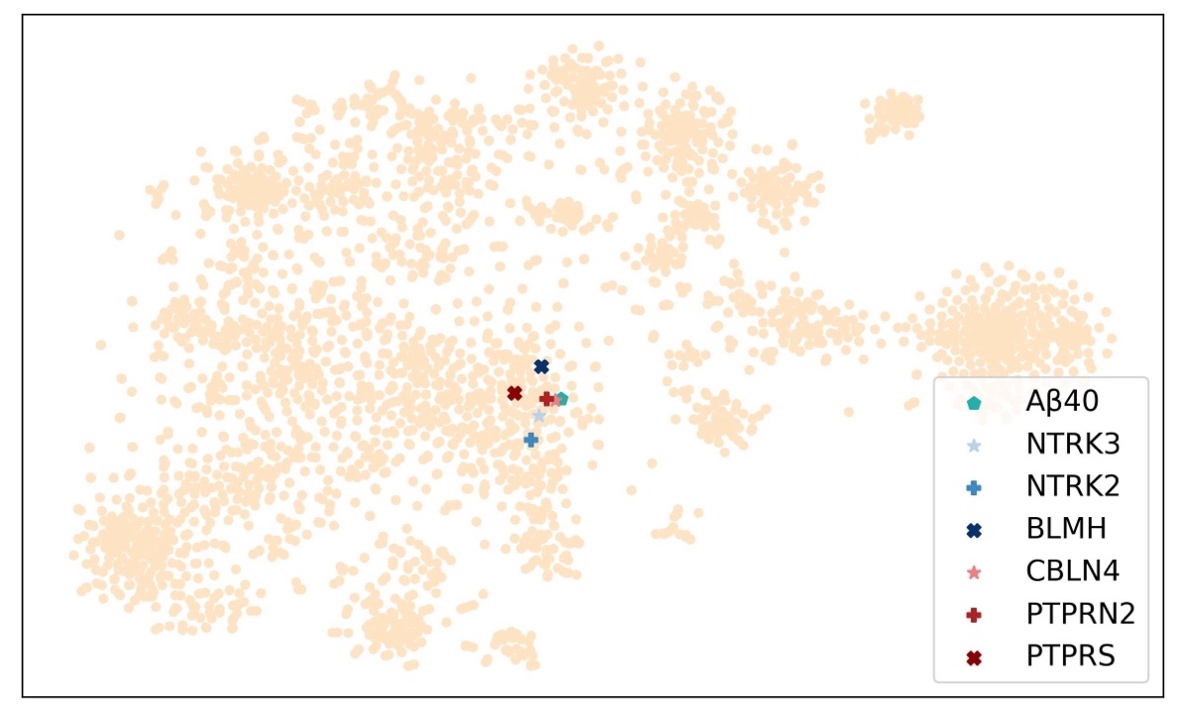

**Supplementary Figure 13: Location of the examined reference protein candidates in t-SNE map.** The examined reference protein candidates in t-SNE space. As seen, the area of interest is covered by single clusters in all examples in Supplementary Fig. 6, indicating that the clustering could be performed with different Ks and random initialization seeds and still include these candidates.

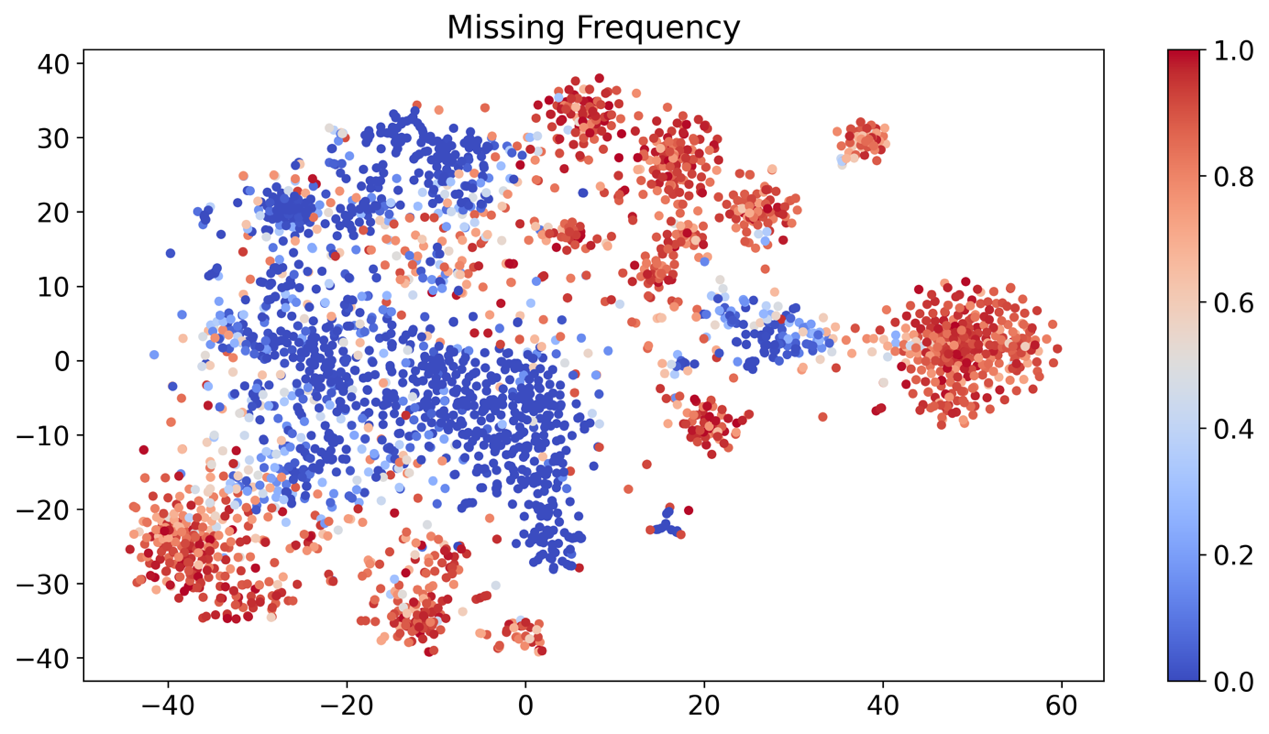

**Supplementary Figure 14: Missing frequency t-SNE map.** t-SNE map colored according to missing frequency for each protein.

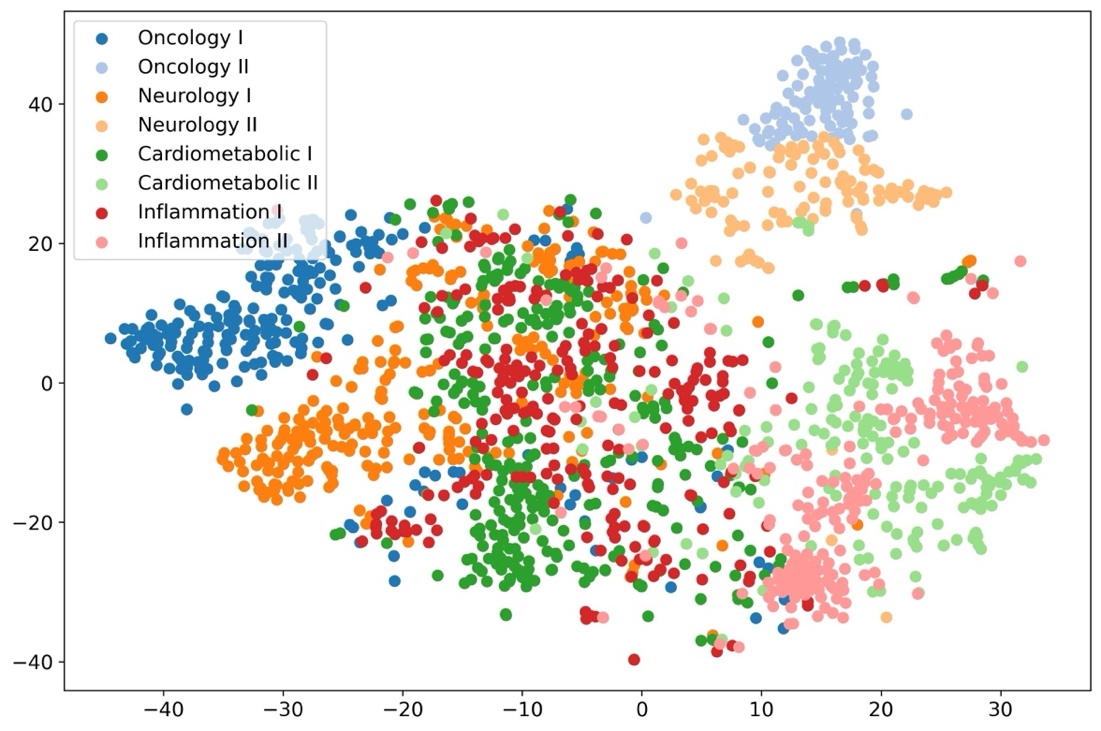

**Supplementary Figure 15: t-SNE on reduced data colored by panel.** t-SNE dimensionality reduction with proteins of missing frequency < 75%, colored by panel. As seen, the panel division is like that of the full dataset (see Supplementary Fig. 4), where Oncology II and Neurology II are separated but close to each other, majority of Oncology I and Neurology I next to each other, and Cardiometabolic II and Inflammation II together.

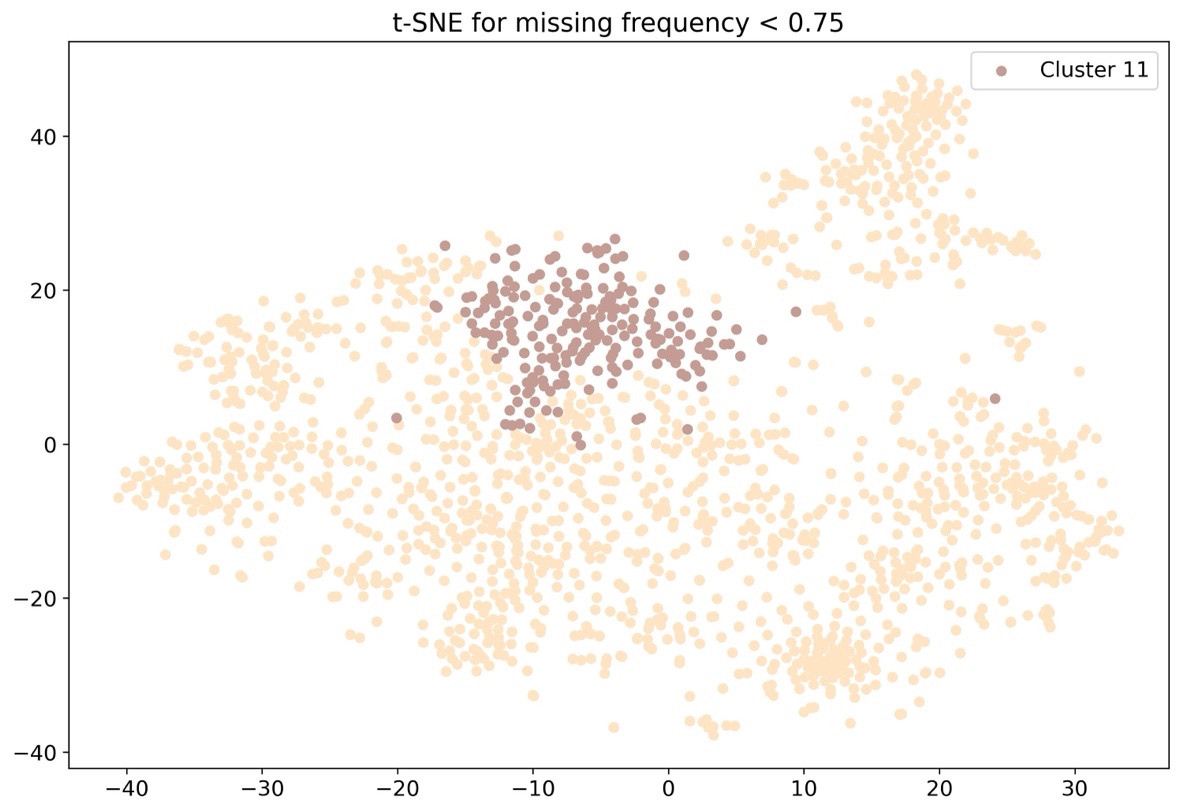

**Supplementary Figure 16: t-SNE on reduced data highlighting cluster 11.** t-SNE dimensionality reduction with proteins of missing frequency <75% highlighting cluster 11. The proteins of the cluster are still located close to each other, covering a central area of the map.

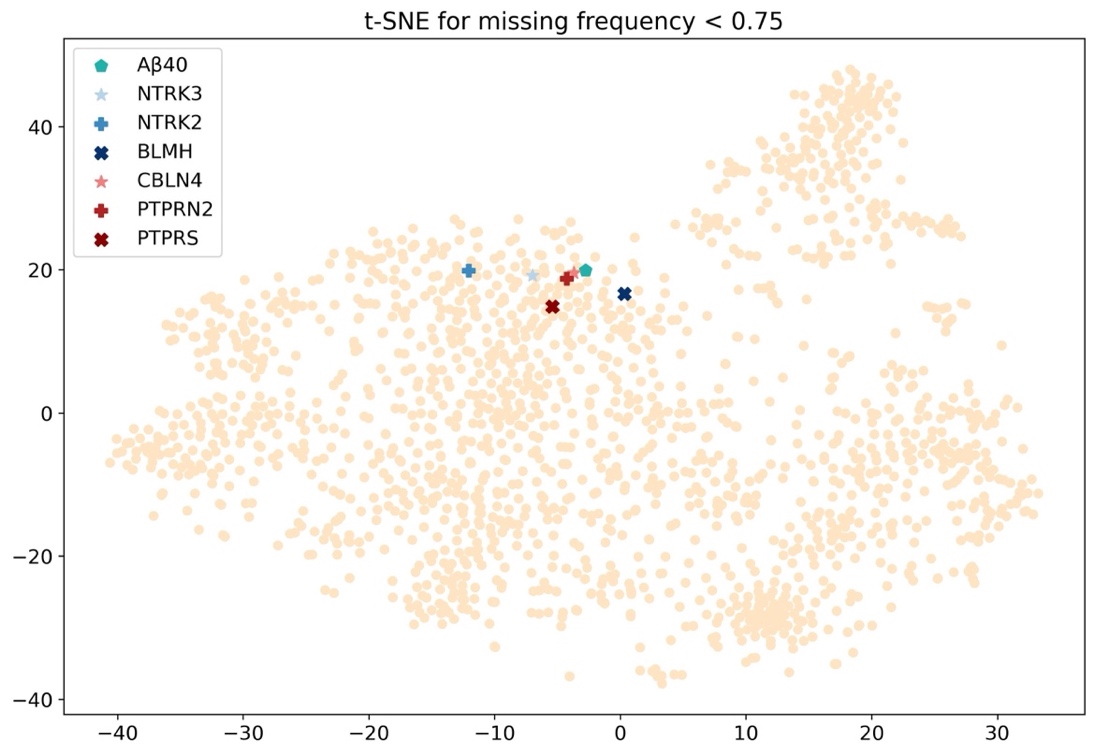

**Supplementary Figure 17: t-SNE on reduced data highlighting reference protein candidates.** t-SNE dimensionality reduction with proteins of missing frequency <75% highlighting the suggested reference protein candidates. For this reduced dataset, the candidates are still located close to each other inside cluster 11.

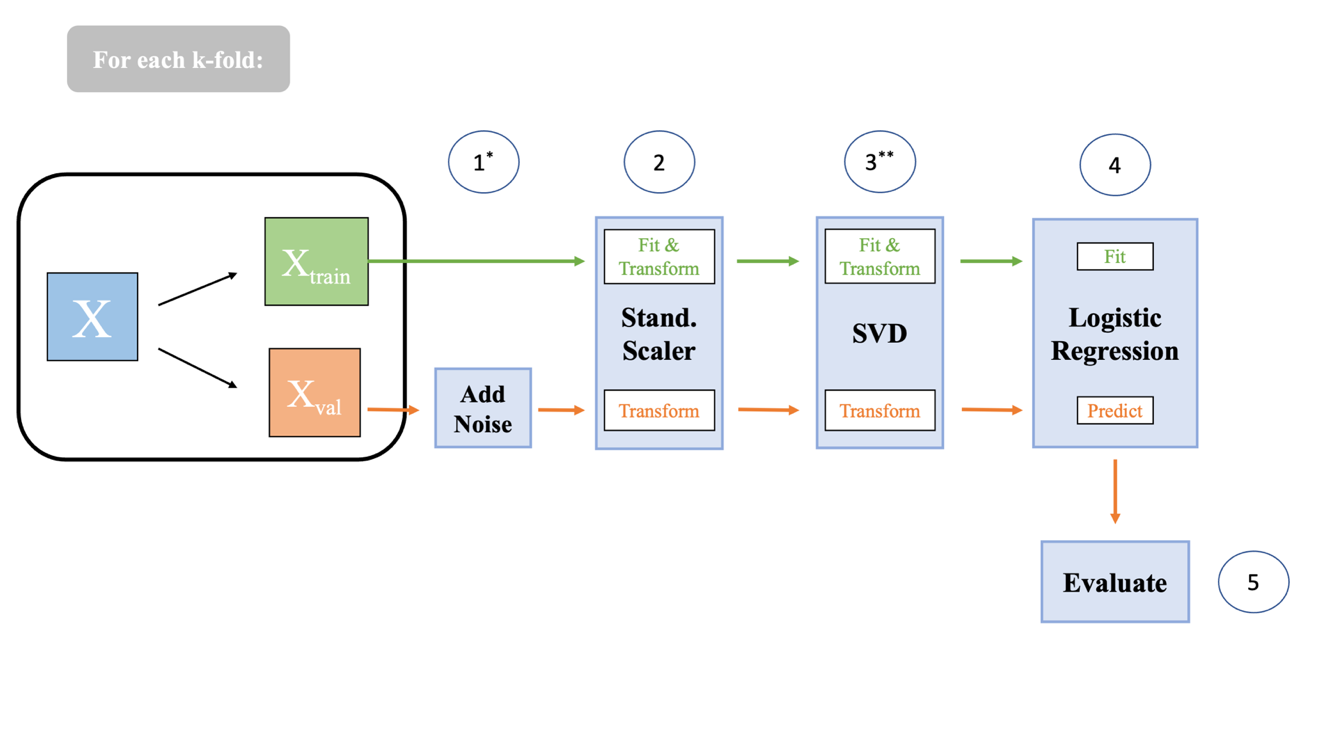

**Supplementary Figure 18**: **Pipeline for pre-processing steps of all models.** All fits (standard scaling, SVD and logistic regression) were applied only to training data (either training dataset or training folds/samples in 10-fold-cross-validation/bootstrap). Both training and validation data were then transformed accordingly. Test data was treated as validation data in this pipeline. Steps 1 and 3 were skipped unless specified otherwise.
* Noise was only added in the simulated noise analysis, see *Supplementary Results: Simulated Measurement Noise Analysis*.
** SVD was only applied when a singular value decomposition was used as reference.
