## Supplementary Tables for "Cerebrospinal fluid reference proteins increase accuracy and interpretability of biomarkers for brain diseases"

**Supplementary Table 1: Predicting the mean CSF protein level in a multiple linear regression model.** Testing the association of independent variables age, sex, education years, intracranial volume, total gray matter volume and ventricular volume with the mean CSF protein level in a multiple linear regression model, with either all participants or cognitively unimpaired Aβ-negative participants only. Significant variables are denoted as bold.

|  | **Age** | **Sex** | **Education** | **Intracranial volume** | **Total gray matter volume** | **Ventricular volume** |
| --- | --- | --- | --- | --- | --- | --- |
| **All Participants (n=631)** *R-squared = 0.256* | | | | | | |
| *β-coef* | **0.544** | **-0.159** | -0.018 | 0.037 | 0.018 | **-0.321** |
| *P-value* | **4e-31** | **2e-4** | 0.6 | 0.6 | 0.8 | **5e-11** |
| **Cognitively unimpaired Aβ-negative participants (n=197)** *R-squared =0.371* | | | | | | |
| *β-coef* | **0.705** | **-0.212** | 0.104 | 0.098 | -0.029 | **-0.348** |
| *P-value* | **3e-13** | **0.002** | 0.08 | 0.4 | 0.8 | **6e-5** |

**Supplementary Table 2:** **Performance details for all models with and without references.** Quantitative details for the three models evaluated on BF2 test dataset and independent cohort BF1, as presented in Fig. 5. **P* < 0.05, ***P* < 0.01

|  | **AUC (95% CI)** | **AUC difference vs. no reference (*P*-value)** | **AUC difference vs. mean CSF (*P*-value)** |
| --- | --- | --- | --- |
| **P-tau181→TauPET (BF2)** | | | |
| No reference | **0.828** (0.750, 0.906) | **-** | **-** |
| Aβ40 | **0.933** (0.896, 0.970) | **0.105** (0.002**) | **0.0386** (0.01*) |
| Mean CSF level | **0.895** (0.847, 0.943) | **0.0663** (0.009**) | **-** |
| NTRK3 | **0.920** (0.879, 0.962) | **0.0981** (0.003**) | **0.0255** (0.01*) |
| NTRK2 | **0.923** (0.882, 0.964) | **0.0947** (0.002**) | **0.0284** (0.01*) |
| BLMH | **0.908** (0.861, 0.954) | **0.0794** (0.005**) | **0.0131** (0.2) |
| SVD1 (NTRK3, NTRK2 and BLMH) | **0.928** (0.889, 0.967) | **0.0995** (0.002**) | **0.0332** (0.01*) |
| CBLN4 | **0.944** (0.907, 0.981) | **0.115** (0.002**) | **0.0491** (0.01*) |
| PTPRN2 | **0.937** (0.901, 0.974) | **0.109** (0.002**) | **0.0427** (0.01*) |
| PTPRS | **0.928** (0.889, 0.967) | **0.0997** (0.003**) | **0.0334** (0.01*) |
| SVD2 (CBLN4, PTPRN2 and PTPRS) | **0.946** (0.913, 0.980) | **0.118** (0.002**) | **0.0518** (0.009**) |
| **Aβ42→AβPET (BF2)** | | | |
| Baseline | **0.966** (0.939, 0.993) | **-** | **-** |
| Aβ40 | **0.992** (0.982, 1) | **0.0261** (0.03*) | **0.0148** (0.1) |
| Mean CSF level | **0.977** (0.956, 0.998) | **0.0113** (0.04*) | **-** |
| NTRK3 | **0.983** (0.966, 1) | **0.0169** (0.03*) | **0.00563** (0.2) |
| NTRK2 | **0.987** (0.972, 1) | **0.0208** (0.03*) | **0.00951** (0.1) |
| BLMH | **0.986** (0.971, 1) | **0.0201** (0.03*) | **0.00880** (0.1) |
| SVD1 | **0.987** (0.972, 1) | **0.0208** (0.03*) | **0.00951** (0.1) |
| CBLN4 | **0.983** (0.966,1) | **0.0176** (0.03*) | **0.00634** (0.2) |
| PTPRN2 | **0.982** (0.965,1) | **0.0162** (0.03*) | **0.00493** (0.2) |
| PTPRS | **0.983** (0.967,1) | **0.0176** (0.03*) | **0.00634** (0.1) |
| SVD2 (CBLN4, PTPRN2 and PTPRS) | **0.985** (0.969, 1) | **0.0190** (0.03*) | **0.00775** (0.1) |
| **P-tau181→ADDconv (BF2)** | | | |
| No reference | **0.866** (0.767,0.950) | **-** | **-** |
| Aβ40 | **0.898** (0.814, 0.965) | **0.0324** (0.2) | **-0.00629** (0.5) |
| Mean CSF level | **0.905** (0.821, 0.970) | **0.0394** (0.1) | **-** |
| NTRK3 | **0.920** (0.856, 0.971) | **0.0544** (0.1) | **0.0157** (0.4) |
| NTRK2 | **0.912** (0.842, 0.967) | **0.0459** (0.2) | **0.00722** (0.4) |
| BLMH | **0.911** (0.843, 0.963) | **0.0447** (0.1) | **0.00595** (0.4) |
| SVD1 (NTRK3, NTRK2 and BLMH) | **0.918** (0.855,0.970) | **0.0524** (0.1) | **0.0142** (0.4) |
| CBLN4 | **0.925** (0.866, 0.971) | **0.0595** (0.1) | **0.0208** (0.4) |
| PTPRN2 | **0.925** (0.866, 0.974) | **0.0593** (0.1) | **0.0206** (0.4) |
| PTPRS | **0.922** (0.859, 0.975) | **0.0561** (0.1) | **0.0173** (0.4) |
| SVD2 (CBLN4, PTPRN2 and PTPRS) | **0.930** (0.870, 0.976) | **0.0639** (0.1) | **0.0258** (0.4) |
| **P-tau181→ADDconv (BF1)** | | | |
| No reference | **0.880** (0.838, 0.920) | **-** | **-** |
| Aβ40 | **0.933** (0.900, 0.962) | **0.0538** (0.02*) | **0.0259** (0.1) |
| Mean CSF level | **0.908** (0.870, 0.946) | **0.0278** (0.06) | **-** |
| NTRK3 | **0.935** (0.906, 0.961) | **0.0546** (0.009**) | **0.266** (0.03*) |
| NTRK2 | **0.932** (0.901, 0.960) | **0.0523** (0.009**) | **0.0248** (0.03*) |
| BLMH | **0.896** (0.854, 0.937) | **0.0169** (0.1) | **-0.0112** (0.8) |
| SVD1 | **0.926** (0.888, 0.958) | **0.0500** (0.03*) | **0.0185** (0.1) |
| **Aβ42→AβPET (BF1)** | | | |
| No reference | **0.916** (0.867, 0.962) | **-** | **-** |
| Aβ40 | **0.970** (0.938, 0.996) | **0.0538** (0.006**) | **0.0455** (0.02*) |
| Mean CSF level | **0.924** (0.877, 0.968) | **0.00879** (0.2) | **-** |
| NTRK3 | **0.932** (885, 0.974) | **0.0162** (0.2) | **0.00791** (0.2) |
| NTRK2 | **0.931** (0.884, 0.971) | **0.0154** (0.2) | **0.00713** (0.2) |
| BLMH | **0.920** (0.870, 0.965) | **0.00458** (0.3) | **-0.00371** (0.6) |
| SVD1 | **0.933** (0.883, 0.975) | **0.0167** (0.2) | **0.00820** (0.2) |

**Supplementary Table 3: Effect sizes of association with CSF P-tau181 for sTREM2, sAXL, sTyro3 and YKL-40 without and with reference proteins.** For each model, the protein is evaluated by β-coef, P-value and R-squared. As seen, all proteins are highly associated with CSF P-tau181 without using a reference, but when adding a reference, the effect size decreased severely/disappeared. All models included NC (n=172), SCD (n=92) and MCI (n=145) participants from BF2 and were adjusted for age and sex.

|  | | **Without reference** | **With reference Aβ40** | | | **With reference NTRK3** | | | | **With reference CBLN4** | |
| --- | --- | --- | --- | --- | --- | --- | --- | --- | --- | --- | --- |
|  |  | **Protein** | **Protein** | | **Aβ40** | **Protein** | | | **NTRK3** | **Protein** | **CBLN4** |
| **Protein:**  **sTREM2** | *β-coef* | 0.346 | 0.102 | | 0.569 | 0.0658 | | | 0.495 | 0.131 | 0.480 |
|  | *P-value* | 1e-11 | 0.03 | | 5e-33 | 0.1 | | | 6e-22 | 6e-3 | 9e-26 |
|  | *R-squared* | 0.20 | 0.44 | | | 0.37 | | | | 0.40 | |
| **Protein:**  **sAXL** | *β-coef* | 0.488 | 0.174 | 0.476 | | 0.176 | | 0.385 | | 0.186 | 0.394 |
|  | *P-value* | 1e-26 | 7e-4 | 2e-19 | | 0.01 | | 7e-8 | | 2e-3 | 4e-11 |
|  | *R-squared* | 0.33 | 0.45 | | | 0.37 | | | | 0.40 | |
| **Protein:**  **sTyro3** | *β-coef* | 0.47 | 0.113 | 0.515 | | 0.0307 | | 0.502 | | 0.136 | 0.435 |
|  | *P-value* | 4e-23 | 0.03 | 2e-21 | | 0.7 | | 4e-10 | | 0.02 | 9e-14 |
|  | *R-squared* | 0.30 | 0.44 | | | 0.36 | | | | 0.39 | |
| **Protein:**  **YKL-40** | *β-coef* | 0.567 | 0.329 | 0.470 | | 0.350 | 0.384 | | | 0.350 | 0.398 |
|  | *P-value* | 5e-26 | 8e-11 | 3e-27 | | 1e-10 | 2e-16 | | | 2e-11 | 1e-19 |
|  | *R-squared* | 0.32 | 0.49 | | | 0.43 | | | | 0.45 | |

**Supplementary Table 4: Effect sizes of association with CSF P-tau181 α-synuclein without and with reference proteins.** The protein is evaluated by β-coef, P-value and R-squared. As seen, α-synuclein is highly associated with CSF P-tau181 without using a reference, but when adding a reference in the model the significance decreases. All models included AD dementia participants (n=210) from BF2 and were adjusted for age and sex.

|  | | **Without reference** | **With reference Aβ40** | | **With reference NTRK3** | | **With reference CBLN4** | |
| --- | --- | --- | --- | --- | --- | --- | --- | --- |
|  |  | **Protein** | **Protein** | **Aβ40** | **Protein** | **NTRK3** | **Protein** | **CBLN4** |
| **Protein:**  **α-synuclein** | *β-coef* | 0.37 | 0.213 | 0.589 | 0.222 | 0.484 | 0.207 | 0.541 |
|  | *P-value* | 8e-8 | 2e-4 | 3e-20 | 3e-4 | 1e-12 | 4e-4 | 9e-17 |
|  | *R-squared* | 0.19 | 0.48 | | 0.38 | | 0.44 | |

**Supplementary Table 5: Effect sizes of *APOE*** **ε4 allele association with ApoE4 CSF protein expression without and with reference proteins.** The protein is evaluated by β-coef, P-value and R-squared. The alleles were encoded as follows: ε2ε4, ε3ε4 = 1 and ε4ε4=2. The BF1 dataset was used, only including participants with one or two ε4 alleles and complete measures for all other parameters (n=437).

|  | | **Without reference** | | **With reference Aβ40** | | **With reference NTRK3** | |
| --- | --- | --- | --- | --- | --- | --- | --- |
|  |  | ***APOE* genotype** | ***APOE* genotype** | | **Aβ40** | ***APOE* genotype** | **NTRK3** |
| **Outcome Protein: ApoE4** | *β-coef* | 0.580 | 0.590 | | 0.429 | 0.578 | 0.441 |
|  | *P-value* | 3e-40 | 2e-45 | | 1e-27 | 1e-51 | 4e-33 |
|  | *R-squared* | 0.34 | | 0.52 | | 0.53 | |

**Supplementary Table 6: The selected pQTL-analysis proteins and their association with the mean CSF level.** Linear regression models adjusted for age and sex, describing each protein’s association with the mean CSF level in the BF1 dataset. The higher the association, the more likely to benefit from adjusting for the individual reference level when used as a biomarker.

| **Protein** | **Linear Regression Parameters** |
| --- | --- |
| *VEGFA* | β = 0.82, p<1e-120 |
| *sVCAM1* | β = 0.64, p<1e-40 |
| *PLXNB1* | β = 0.82, p<1e-180 |
| *PRTG* | β = 0.85, p<1e-210 |
| *TFF3* | β = 0.77, p<1e-120 |

**Supplementary Table 7: Effect of genotype (SNP) in associations with CSF proteins without and with reference proteins.** The SNP and reference proteins are evaluated by β-coefficient and P-value, and each model’s R-squared is given. In general, the trans-pQTL associations of the GMNC-OSTN were severely weakened when adjusting for a reference protein. All models included BF1 participants (n=1445) and were adjusted for age, sex, dementia diagnosis and ten genetic principal components.

|  | | **Without reference** | **With reference Aβ40** | | **With reference NTRK3** | |
| --- | --- | --- | --- | --- | --- | --- |
|  |  | **SNP** | **SNP** | **Aβ40** | **SNP** | **NTRK3** |
| **Outcome Protein: VEGFA**  **SNP: rs57712768** | *β-coef* | 0.15 | 0.094 | 0.63 | 0.020 | 0.86 |
|  | *P-value* | 2e-8 | 6e-6 | 1e-145 | 0.1 | <1e-150 |
|  | *R-squared* | 0.089 | 0.45 | | 0.76 | |
| **Outcome Protein: *sVCAM1***  **SNP: rs146550622** | *β-coef* | 0.15 | 0.12 | 0.35 | 0.089 | 0.37 |
|  | *P-value* | 8e-8 | 8e-6 | 1e-36 | 4e-4 | 2e-42 |
|  | *R-squared* | 0.20 | 0.31 | | 0.31 | |
| **Outcome Protein: *PLXNB1***  **SNP: rs4687181** | *β-coef* | 0.16 | 0.080 | 0.71 | 0.027 | 0.86 |
|  | *P-value* | 2e-9 | 5e-5 | 4e-191 | 0.06 | <1e-150 |
|  | *R-squared* | 0.054 | 0.52 | | 0.72 | |
| **Outcome Protein: PRTG**  **SNP: rs71635338** | *β-coef* | 0.16 | 0.11 | 0.52 | 0.022 | 0.89 |
|  | *P-value* | 9e-10 | 3e-6 | 5e-88 | 0.07 | <1e-150 |
|  | *R-squared* | 0.073 | 0.32 | | 0.78 | |
| **Outcome Protein: TFF3**  **SNP: rs78054167** | *β-coef* | 0.15 | 0.11 | 0.33 | 0.039 | 0.62 |
|  | *P-value* | 2e-8 | 4e-6 | 1e-36 | 0.04 | 5e-164 |
|  | *R-squared* | 0.16 | 0.26 | | 0.49 | |

**Supplementary Table 8: OLINK Panel statistics.** Panel statistics of proteins with missing frequency > 50%.

| **OLINK Panel** | **Proteins with Missing Frequency > 50%** |
| --- | --- |
| Explore 384 Cardiometabolic | 19% |
| Explore 384 Cardiometabolic II | 59% |
| Explore 384 Inflammation | 43% |
| Explore 384 Inflammation II | 42% |
| Explore 384 Neurology | 35% |
| Explore 384 Neurology II | 78% |
| Explore 384 Oncology | 45% |
| Explore 384 Oncology II | 84% |

**Supplementary Table 9: Association with the mean CSF level for the proteins that were analyzed.** Linear regression models adjusted for age and sex, describing each protein’s association with the mean CSF level in BF2 training dataset. Sorted in decreasing order. The higher the association, the more likely to benefit from adjusting for the individual reference level when used as a biomarker.

| **Protein** | **Linear Regression Result** |
| --- | --- |
| *sAXL* | β = 0.75, p<1e-150 |
| *sTyro3* | β = 0.68, p<1e-100 |
| *sTREM2* | β = 0.51, p<1e-40 |
| *YKL-40* | β = 0.40, p<1e-20 |
| *GFAP* | β = 0.38, p<1e-20 |
| *Neurogranin* | β = 0.35, p<1e-20 |
| *T-tau* | β = 0.34, p<1e-20 |
| *P-tau* | β = 0.34, p<1e-20 |
| *S100* | β = 0.29, p<1e-10 |
| *Aβ42* | β = 0.24, p<1e-10 |
| *Alpha Synuclein* | β = 0.23, p<1e-10 |
| *NfL* | β = 0.10, p<0.007 |
