## Supplementary Reference Protein Candidates for "Cerebrospinal fluid reference proteins increase accuracy and interpretability of biomarkers for brain diseases"

### Reference Protein Candidate Profiles

This section describes the seven reference protein candidates (NTRK3, NTRK2, BLMH, CBLN4, PTPRN2, PTPRS and Aβ40) from a biological perspective. This analysis is fully based on protein descriptions in the Human Protein Atlas^1–3^ (website: [www.proteinatlas.org](http://www.proteinatlas.org)).

#### Neurotrophic receptor tyrosine kinase 3 (NTRK3)

NTRK3 is a membrane-bound receptor that phosphorylates itself and members of the MAPK pathway after neurotrophin binding. Its signaling controls cell survival and differentiation, mainly in neurons. NTRK3 has enhanced specificity in brain tissue but low regional brain specificity. It is an intracellular and membrane bound protein, expressed in the nucleus and cytoplasm. Data characteristics in BF2 can be seen in Supplementary Candidate Fig. 1, 2 and Tab. 1.

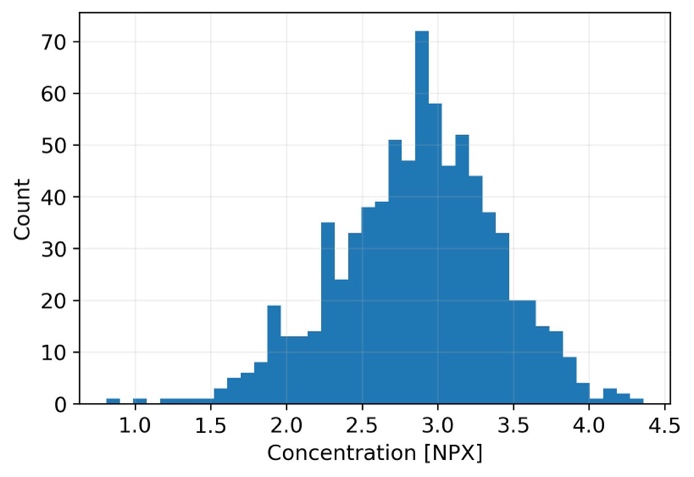

**Supplementary Candidate Figure 1: NTRK3 concentration distribution.** Distribution of protein concentration in BF2 training data.

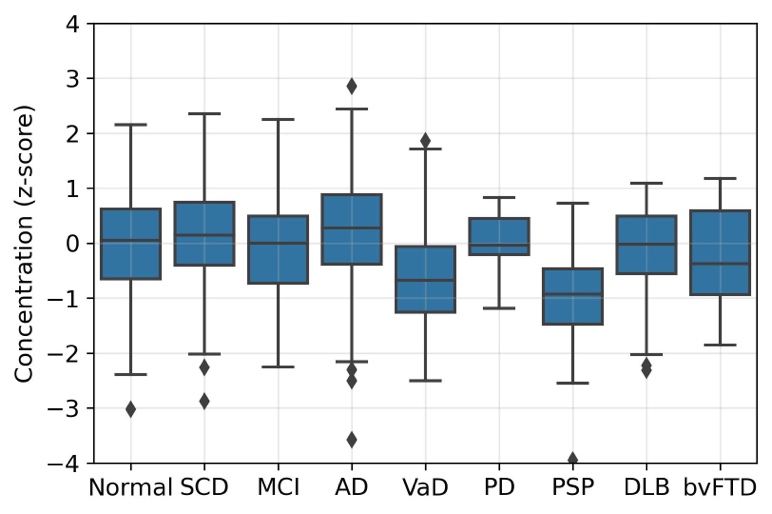

**Supplementary Candidate Figure 2: NTRK3 diagnostic group boxplot.** Distribution of standardized protein concentration between diagnostic groups (>10 individuals) for BF2 training data. Number of participants in each group: normal cognition: n = 196, subjective cognitive decline (SCD): n = 98, mild cognitive impairment (MCI): n = 50, Alzheimer’s disease (AD): n = 237, vascular dementia (VaD): n = 26, Parkinson’s disease (PD): n = 12, Progressive supranuclear palsy (PSP): n = 17, dementia with Lewy bodies (DLB): n = 30, behavioral variant of Frontotemporal dementia (bvFTD): n = 21.

**Supplementary Candidate Table 1: NTRK3 protein data info.** Analysis info from OLINK measures. Association (from linear regression model) with the main predictor to estimate suitability as reference protein for this biomarker. AUC without main predictor to estimate reference candidate’s predictive power without main biomarker. All adjusted for age and sex. An optimal reference should be associated with the main predictor while not being highly predictive of the outcome without the main predictor.

| **Association with mean CSF level** | |
| --- | --- |
| **Linear regression** | β = 0.73, p<1e-130 |
| **Partial correlation (Pearson)** | 0.78 |
| **OLINK analysis info** | |
| Panel | Neurology |
| Mean LOD [NPX] | -5.0 |
| Missing Frequency [%] | 0 |
| **CSF P-tau181 🡪 Tau PET** | |
| Association with main predictor | β = 0.48, p<1e-40 |
| Mean AUC without main predictor | 0.62 |
| **CSF Aβ42 🡪 Aβ PET** | |
| Association with main predictor | β = 0.44, p<1e-30 |
| Mean AUC without main predictor | 0.65 |

#### Neurotrophic receptor tyrosine kinase 2 (NTRK2)

NTRK2 is a membrane-bound receptor that phosphorylates itself and members of the MAPK pathway after neurotrophin binding. Its signaling controls cell survival, proliferation, migration, synapse formation and differentiation, mainly in neurons. NTRK3 has enhanced specificity in brain (and thyroid gland) tissue but low regional brain specificity. It is a membrane bound protein, expressed in the cytoplasm. Data characteristics in BF2 can be seen in Supplementary Candidate Fig. 3, 4 and Tab. 2.

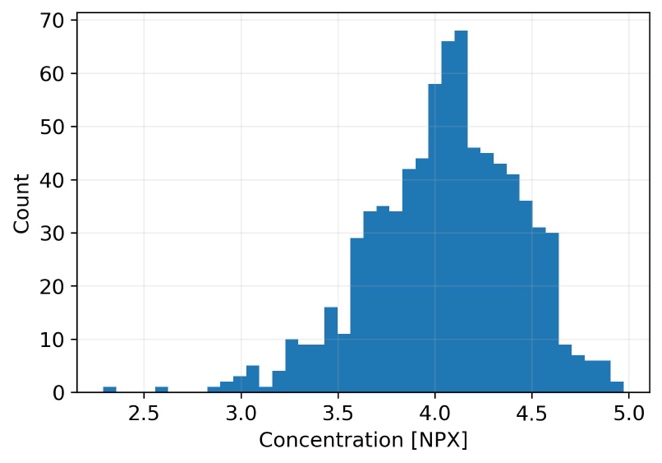

**Supplementary Candidate Figure 3: NTRK2 concentration distribution.** Distribution of protein concentration in BF2 training data.

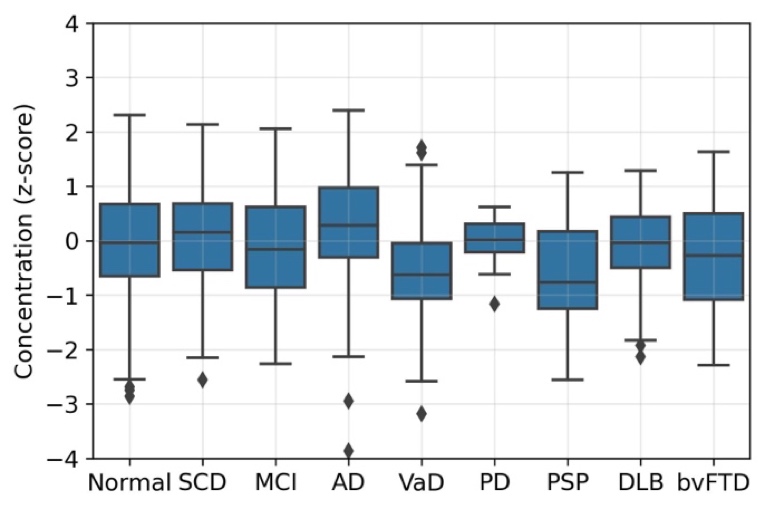

**Supplementary Candidate Figure 4: NTRK2 diagnostic group boxplot.** Distribution of standardized protein concentration between diagnostic groups (>10 individuals) for BF2 training data. Number of participants in each group: normal cognition: n = 196, subjective cognitive decline (SCD): n = 98, mild cognitive impairment (MCI): n = 50, Alzheimer’s disease (AD): n = 237, vascular dementia (VaD): n = 26, Parkinson’s disease (PD): n = 12, Progressive supranuclear palsy (PSP): n = 17, dementia with Lewy bodies (DLB): n = 30, behavioral variant of Frontotemporal dementia (bvFTD): n = 21.

**Supplementary Candidate Table 2: NTRK2 protein data info.** Analysis info from OLINK measures. Association (from linear regression model) with the main predictor to estimate suitability as reference protein for this biomarker. AUC without main predictor to estimate reference candidate’s predictive power without main biomarker. All adjusted for age and sex. An optimal reference should be associated with the main predictor while not being highly predictive of the outcome without the main predictor.

| **Association with mean CSF level** | |
| --- | --- |
| **Linear regression** | β = 0.66, p<1e-100 |
| **Partial correlation (Pearson)** | 0.71 |
| **OLINK analysis info** | |
| Panel | Cardiometabolic |
| Mean LOD [NPX] | -2.6 |
| Missing Frequency [%] | 0 |
| **CSF P-tau181 🡪 Tau PET** | |
| Association with main predictor | β = 0.50, p<1e-40 |
| Mean AUC without main predictor | 0.62 |
| **CSF Aβ42 🡪 Aβ PET** | |
| Association with main predictor | β = 0.40, p<1e-20 |
| Mean AUC without main predictor | 0.66 |

#### Bleomycin hydrolase (BLMH)

BLMH is a cytoplasmic cysteine peptidase that is highly conserved through evolution. Its normal physiological role is unknown. BLMH has enhanced specificity in skin tissue and has low regional brain specificity. It is an intracellular protein expressed in the cytoplasm. Data characteristics in BF2 can be seen in Supplementary Candidate Fig. 5, 6 and Tab. 3.

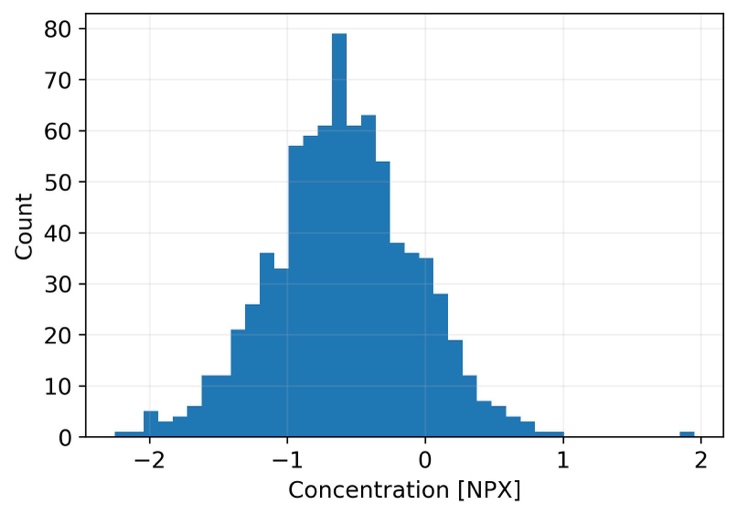

**Supplementary Candidate Figure 5: BLMH concentration distribution.** Distribution of protein concentration in BF2 training data.

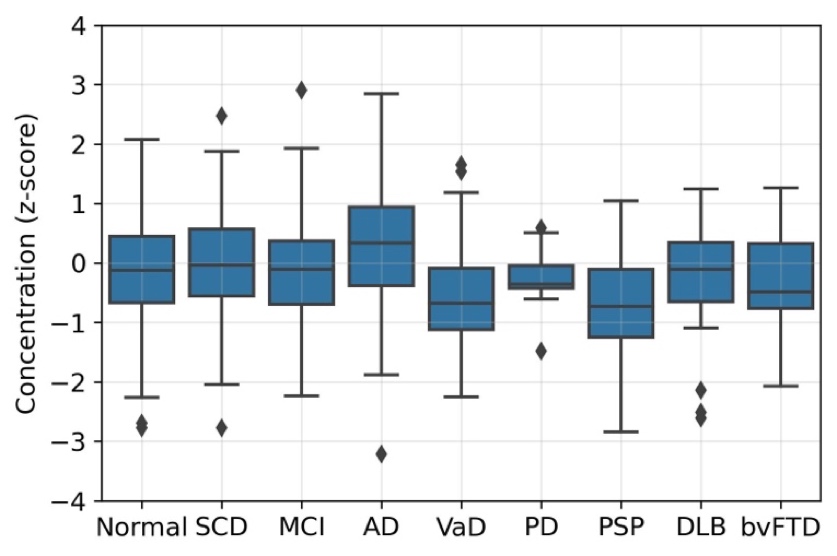

**Supplementary Candidate Figure 6: BLMH diagnostic group boxplot.** Distribution of standardized protein concentration between diagnostic groups (>10 individuals) for BF2 training data. Number of participants in each group: normal cognition: n = 196, subjective cognitive decline (SCD): n = 98, mild cognitive impairment (MCI): n = 50, Alzheimer’s disease (AD): n = 237, vascular dementia (VaD): n = 26, Parkinson’s disease (PD): n = 12, Progressive supranuclear palsy (PSP): n = 17, dementia with Lewy bodies (DLB): n = 30, behavioral variant of Frontotemporal dementia (bvFTD): n = 21.

**Supplementary Candidate Table 3: BLMH protein data info.** Analysis info from OLINK measures. Association (from linear regression model) with the main predictor to estimate suitability as reference protein for this biomarker. AUC without main predictor to estimate reference candidate’s predictive power without main biomarker. All adjusted for age and sex. An optimal reference should be associated with the main predictor while not being highly predictive of the outcome without the main predictor.

| **Association with mean CSF level** | |
| --- | --- |
| **Linear regression** | β = 0.67, p<1e-100 |
| **Partial correlation (Pearson)** | 0.72 |
| **OLINK analysis info** | |
| Panel | Cardiometabolic |
| Mean LOD [NPX] | -6.2 |
| Missing Frequency [%] | 0 |
| **CSF P-tau181 🡪 Tau PET** | |
| Association with main predictor | β = 0.56, p<1e-60 |
| Mean AUC without main predictor | 0.64 |
| **CSF Aβ42 🡪 Aβ PET** | |
| Association with main predictor | β = 0.33, p<1e-15 |
| Mean AUC without main predictor | 0.67 |

#### Cerebellin 4 precursor (CBLN4)

CBLN4 is a synaptic organizer that is involved in regulation of neurexin signaling during synapse development. The protein has enhanced specificity in adrenal gland, brain and epididymis tissue. It has low regional brain specificity. Cellular location mainly in synapses in neurons, and extracellularly secreted in brain. Data characteristics in BF2 can be seen in Supplementary Candidate Fig. 7, 8 and Tab. 4.

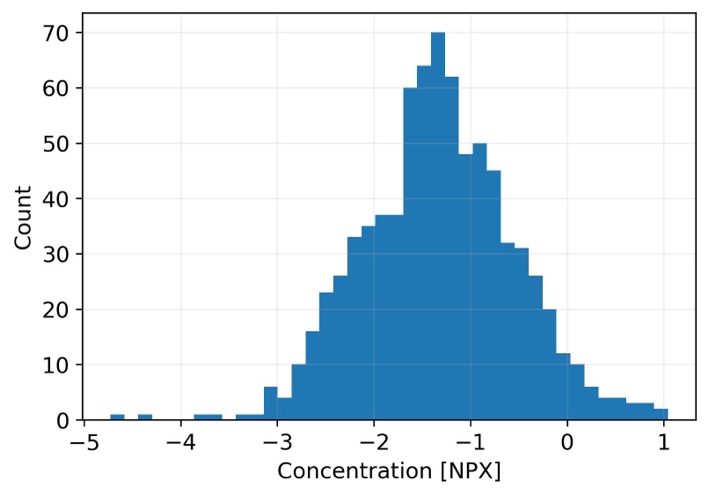

**Supplementary Candidate Figure 7: CBLN4 concentration distribution.** Distribution of protein concentration in BF2 training data.

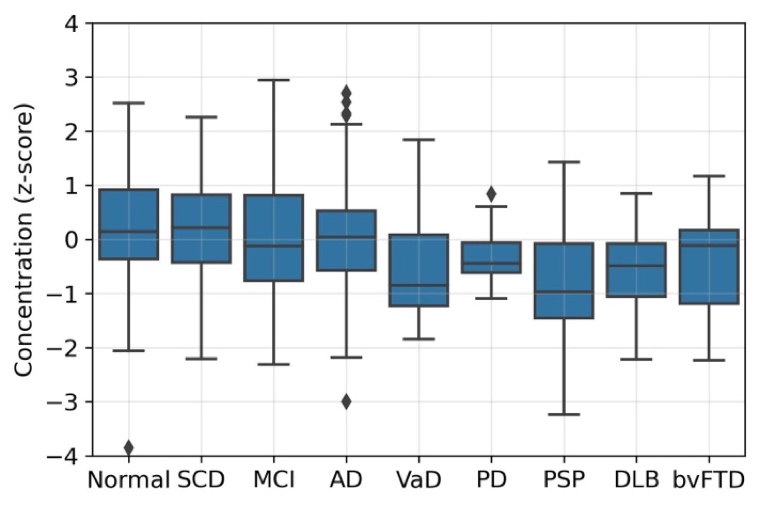

**Supplementary Candidate Figure 8: CBLN4 diagnostic group boxplot.** Distribution of standardized protein concentration between diagnostic groups (>10 individuals) for BF2 training data. Number of participants in each group: normal cognition: n = 196, subjective cognitive decline (SCD): n = 98, mild cognitive impairment (MCI): n = 50, Alzheimer’s disease (AD): n = 237, vascular dementia (VaD): n = 26, Parkinson’s disease (PD): n = 12, Progressive supranuclear palsy (PSP): n = 17, dementia with Lewy bodies (DLB): n = 30, behavioral variant of Frontotemporal dementia (bvFTD): n = 21.

**Supplementary Candidate Table 4: CBLN4 protein data info.** Analysis info from OLINK measures. Association (from linear regression model) with the main predictor to estimate suitability as reference protein for this biomarker. AUC without main predictor to estimate reference candidate’s predictive power without main biomarker. All adjusted for age and sex. An optimal reference should be associated with the main predictor while not being highly predictive of the outcome without the main predictor.

| **Association with mean CSF level** | |
| --- | --- |
| **Linear regression** | β = 0.56, p<1e-70 |
| **Partial correlation (Pearson)** | 0.62 |
| **OLINK analysis info** | |
| Panel | Oncology |
| Mean LOD [NPX] | -4.5 |
| Missing Frequency [%] | 0.0008 |
| **CSF P-tau181 🡪 Tau PET** | |
| Association with main predictor | β = 0.44, p<1e-30 |
| Mean AUC without main predictor | 0.64 |
| **CSF Aβ42 🡪 Aβ PET** | |
| Association with main predictor | β = 0.47, p<1e-40 |
| Mean AUC without main predictor | 0.65 |

#### Protein tyrosine phosphatase receptor type N2 (PTPRN2)

PTPRN2 is involved in the vesicle-mediated secretory processes, where it is required for normal accumulation of secretory vesicles in hippocampus, pituitary and pancreatic islets. The protein has enhanced specificity in brain and pancreas tissue. It has low regional brain specificity and is expressed in the brain cells’ cytoplasm. PTPRN2 is an intracellular and membrane bound protein. Data characteristics in BF2 can be seen in Supplementary Candidate Fig. 9, 10 and Tab. 5.

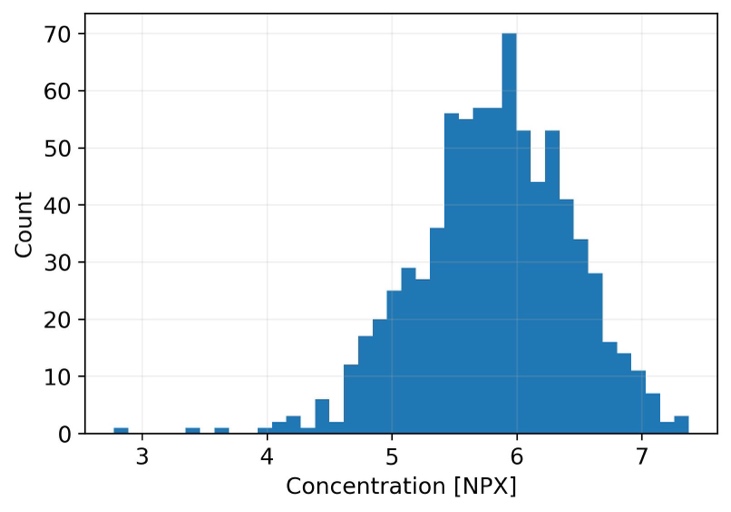

**Supplementary Candidate Figure 9: PTPRN2 concentration distribution.** Distribution of protein concentration in BF2 training data.

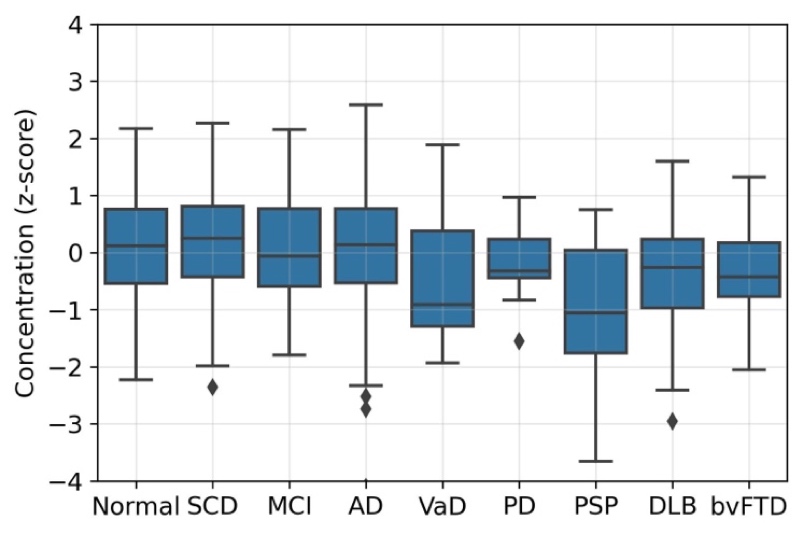

**Supplementary Candidate Figure 10: PTPRN2 diagnostic group boxplot.** Distribution of standardized protein concentration between diagnostic groups (>10 individuals) for BF2 training data. Number of participants in each group: normal cognition: n = 196, subjective cognitive decline (SCD): n = 98, mild cognitive impairment (MCI): n = 50, Alzheimer’s disease (AD): n = 237, vascular dementia (VaD): n = 26, Parkinson’s disease (PD): n = 12, Progressive supranuclear palsy (PSP): n = 17, dementia with Lewy bodies (DLB): n = 30, behavioral variant of Frontotemporal dementia (bvFTD): n = 21.

**Supplementary Table 5: PTPRN2 protein data info.** Analysis info from OLINK measures. Association (from linear regression model) with the main predictor to estimate suitability as reference protein for this biomarker. AUC without main predictor to estimate reference candidate’s predictive power without main biomarker. All adjusted for age and sex. An optimal reference should be associated with the main predictor while not being highly predictive of the outcome without the main predictor.

| **Association with mean CSF level** | |
| --- | --- |
| **Linear regression** | β = 0.65, p<1e-100 |
| **Partial correlation (Pearson)** | 0.71 |
| **OLINK analysis info** | |
| Panel | Neurology |
| Mean LOD [NPX] | -3.6 |
| Missing Frequency [%] | 0 |
| **CSF P-tau181 🡪 Tau PET** | |
| Association with main predictor | β = 0.48, p<1e-40 |
| Mean AUC without main predictor | 0.63 |
| **CSF Aβ42 🡪 Aβ PET** | |
| Association with main predictor | β = 0.49, p<1e-40 |
| Mean AUC without main predictor | 0.65 |

#### Protein tyrosine phosphatase receptor type S (PTPRS)

PTPRS is a cell surface receptor that contributes to the regulation of neurite and axonal outgrowth, differentiation, mitotic cycle and oncogenic transformation. The protein has low tissue specificity and low regional brain specificity. It is mainly expressed in neurons and located in the cytosol and plasma membrane. PTPRS is a membrane bound protein and extracellularly secreted. Data characteristics in BF2 can be seen in Supplementary Candidate Fig. 11, 12 and Tab. 6.

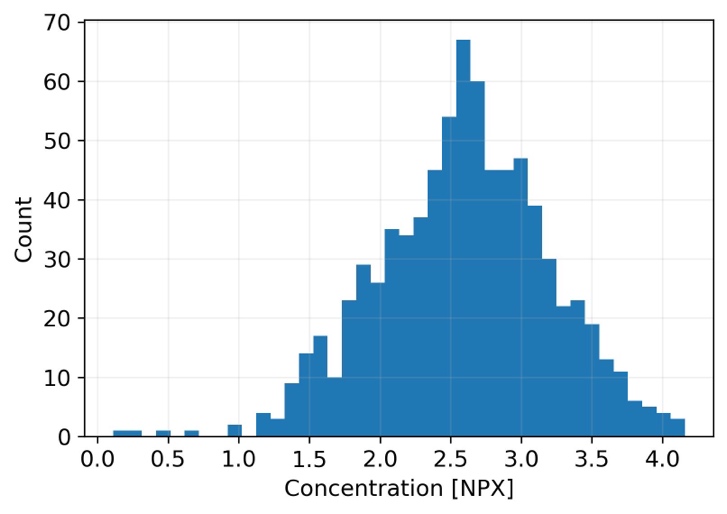

**Supplementary Candidate Figure 11: PTPRS concentration distribution.** Distribution of protein concentration in BF2 training data

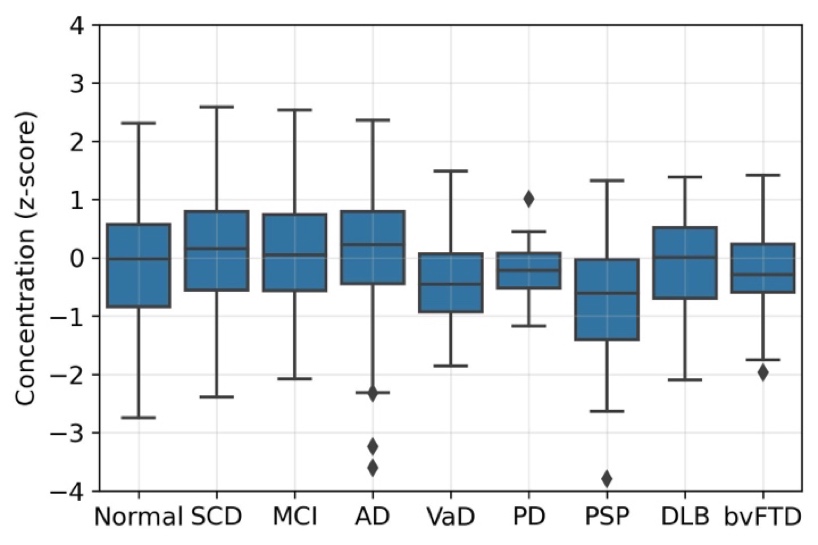

**Supplementary Candidate Figure 12: PTPRS diagnostic group boxplot.** Distribution of standardized protein concentration between diagnostic groups (>10 individuals) for BF2 training data. Number of participants in each group: normal cognition: n = 196, subjective cognitive decline (SCD): n = 98, mild cognitive impairment (MCI): n = 50, Alzheimer’s disease (AD): n = 237, vascular dementia (VaD): n = 26, Parkinson’s disease (PD): n = 12, Progressive supranuclear palsy (PSP): n = 17, dementia with Lewy bodies (DLB): n = 30, behavioral variant of Frontotemporal dementia (bvFTD): n = 21.

**Supplementary Candidate Table 6: PTPRS protein data info.** Analysis info from OLINK measures. Association (from linear regression model) with the main predictor to estimate suitability as reference protein for this biomarker. AUC without main predictor to estimate reference candidate’s predictive power without main biomarker. All adjusted for age and sex. An optimal reference should be associated with the main predictor while not being highly predictive of the outcome without the main predictor.

| **Association with mean CSF level** | |
| --- | --- |
| **Linear regression** | β = 0.76, p<1e-140 |
| **Partial correlation (Pearson)** | 0.80 |
| **OLINK analysis info** | |
| Panel | Neurology |
| Mean LOD [NPX] | -3.6 |
| Missing Frequency [%] | 0 |
| **CSF P-tau181 🡪 Tau PET** | |
| Association with main predictor | β = 0.48, p<1e-40 |
| Mean AUC without main predictor | 0.63 |
| **CSF Aβ42 🡪 Aβ PET** | |
| Association with main predictor | β = 0.49, p<1e-40 |
| Mean AUC without main predictor | 0.65 |

#### Amyloid-β40 (Aβ40)

Aβ40 is a peptide derived from the amyloid beta precursor protein (APP). APP is a cell surface receptor that contributes to neurite growth, neuronal adhesion and axonogenesis. It has low tissue specificity and low regional brain specificity. It is a membrane bound protein, mainly expressed in the cytoplasm and secreted extracellularly. Data characteristics in BF2 can be seen in Supplementary Candidate Fig. 13, 14 and Tab. 7.

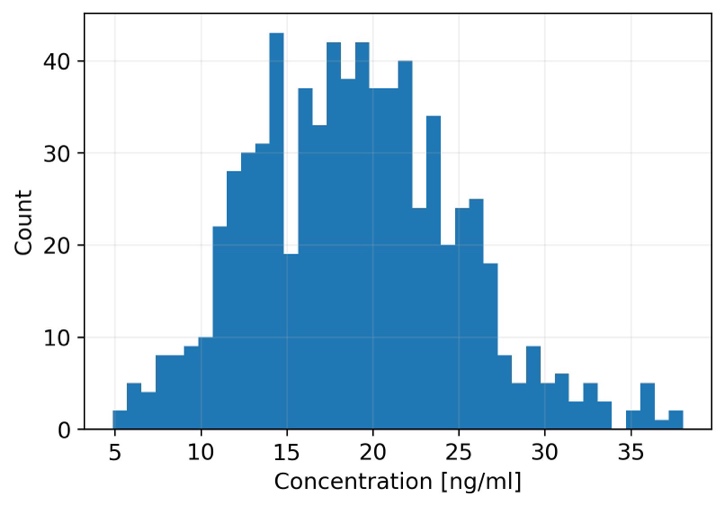

**Supplementary Candidate Figure 13: Aβ40 concentration distribution.** Distribution of protein concentration in BF2 training data.

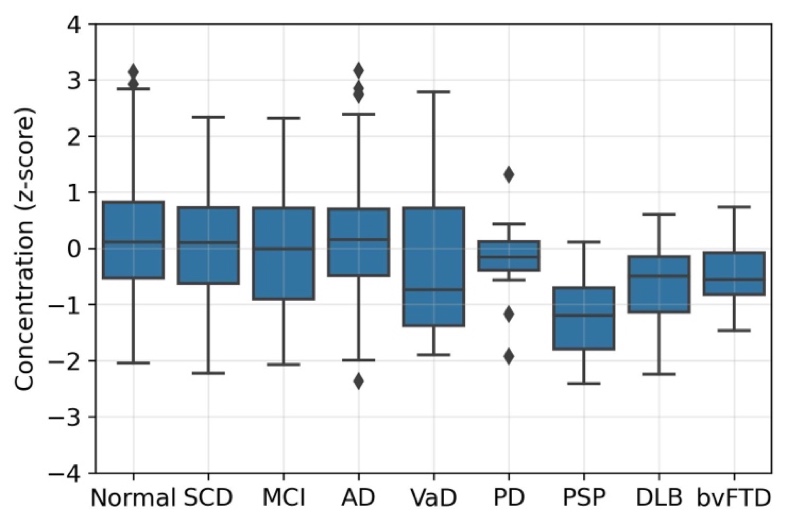

**Supplementary Candidate Figure 14: Aβ40 diagnostic group boxplot.** Distribution of standardized protein concentration between diagnostic groups (>10 individuals) for BF2 training data. Number of participants in each group: normal cognition: n = 196, subjective cognitive decline (SCD): n = 98, mild cognitive impairment (MCI): n = 50, Alzheimer’s disease (AD): n = 237, vascular dementia (VaD): n = 26, Parkinson’s disease (PD): n = 12, Progressive supranuclear palsy (PSP): n = 17, dementia with Lewy bodies (DLB): n = 30, behavioral variant of Frontotemporal dementia (bvFTD): n = 21.

**Supplementary Candidate 7: Aβ40** **protein data info**. This protein was measured with an ELISA assay, therefore no OLINK info exist. Association (from linear regression model) with the main predictor to estimate suitability as reference protein for this biomarker. AUC without main predictor to estimate reference candidate’s predictive power without main biomarker. All adjusted for age and sex. An optimal reference should be associated with the main predictor while not being highly predictive of the outcome without the main predictor.

| **Association with mean CSF level** | |
| --- | --- |
| **Linear regression** | β = 0.44, p<1e-30 |
| **Partial correlation (Pearson)** | 0.48 |
| **CSF P-tau181 🡪 Tau PET** | |
| Association with main predictor | β = 0.50, p<1e-50 |
| Mean AUC without main predictor | 0.62 |
| **CSF Aβ42 🡪 Aβ PET** | |
| Association with main predictor | β = 0.52, p<1e-40 |
| Mean AUC without main predictor | 0.65 |

### References

1.	Uhlén, M. *et al.* Tissue-based map of the human proteome. *Science (1979)* **347**, (2015).

2.	Pontén, F., Jirström, K. & Uhlen, M. The Human Protein Atlas - A tool for pathology. *Journal of Pathology* vol. 216 387–393 Preprint at https://doi.org/10.1002/path.2440 (2008).

3.	Uhlen, M. *et al.* Towards a knowledge-based Human Protein Atlas. *Nature Biotechnology* vol. 28 1248–1250 Preprint at https://doi.org/10.1038/nbt1210-1248 (2010).
