## Supplementary Methods for "Cerebrospinal fluid reference proteins increase accuracy and interpretability of biomarkers for brain diseases"

### OLINK Data Handling

#### Quality Control

Each OLINK sample has gone through three internal and three external quality control steps. The internal controls (Incubation, Extension and Amplification control) are used to monitor quality of assay performance and individual samples, as well as generation of NPX values. The external controls are used for data normalization, to assess variation between runs and plates, to calculate limit of detection (LOD) and discover contamination. All samples are included in the data output file but labeled with WARN if they do not pass a certain quality control. 95% of all datapoints passed QC for BF2. All data was kept for the analysis to maintain statistical power, as the existing QC/assay warnings were distributed between different proteins and participants.

#### Limit of Detection

Limit of Detection (LOD) is calculated from the background plus three standard deviations (assay specific), estimated from negative controls. LODs are computed separately for each assay and plate. Due to the S-curve relationship of OLINK platform data, concentrations below LOD are at risk of being non-linear (meaning that 1 NPX differences do not correspond to 2x change). This may bias results as values tend to be condensed to a small range. Still, the data can contain informative structures and can differ between groups, which is why OLINK do not recommend removing data points below LOD. No LOD filtering was therefore performed in this work. The percentage of proteins below LOD are referred to as the missing frequency. For the BF2 CSF OLINK data, the within-panel statistics of number of proteins with missing frequency > 50% can be seen in Supplementary Tab. 8.

### LOD Sensitivity Analysis

To create Fig. 2 and 3, all 2,943 OLINK proteins were included. To estimate the effect of low LODs, the t-SNE map was colored according to missing frequency in Supplementary Fig. 14. As seen, the area of highly detected proteins (blue) overlaps with the area of proteins strongly associated with the mean CSF level (see Fig. 3c). Additionally, the most well-separated clusters tend to contain a high extent of proteins below LOD. As there still exist clear clustering characteristics for proteins with high missing frequency, both within and between panels (compare with Supplementary Fig. 3), the data evidently contains identifying structures. We therefore considered it reasonable to include all the proteins in this analysis. To ensure these qualities did not affect conclusions about cluster 11, a sensitivity analysis of the visualization of concept and t-SNE dimensionality reduction was performed, filtering out all proteins with missing frequency > 75%. When doing so, 1,730 proteins maintained included. The result can be seen in Supplementary Fig. 15-17 and are in line with the findings from using all 2,943 OLINK proteins.

### K-means Robustness Analysis

The K-means robustness analysis aims to provide evidence that the results were not heavily dependent on the selection of K or random initialization seed. A silhouette plot is provided (Supplementary Fig. 11), showing how selecting a certain K is neither obvious nor of great importance for generating similar results. The silhouette score is a metric that evaluates the quality of a clustering result, considering cluster wise intra- and inter-distances. The score ranges between -1 and 1, where 1 represents a dense and well-separated clustering, 0 overlapping clusters with samples nearby decision boundaries and negative values that samples have likely been assigned to wrong clusters^1^. In addition to the silhouette plot, new versions with different Ks and random initialization seeds of the clustering in the t-SNE map are shown (Supplementary Fig. 12). As seen, similar results as was presented in this work can easily be reproduced, as the nominated reference protein candidates were for all tested Ks and random initializations clustered together (compare with Supplementary Fig. 13).

### References

1.	Shahapure KR, Nicholas C. Cluster quality analysis using silhouette score. In: *Proceedings - 2020 IEEE 7th International Conference on Data Science and Advanced Analytics, DSAA 2020*. Institute of Electrical and Electronics Engineers Inc.; 2020:747-748. doi:10.1109/DSAA49011.2020.00096
